# A multi-tiered μDicer with hierarchical blades achieves protein-preserving microdissection down to 10 μm

**DOI:** 10.64898/2025.12.12.694048

**Authors:** Annatoma Arif, Rashmi Kumar, Yumi Kwon, Ramon Rodriguez, Seth C. Cordts, Saisneha Koppaka, Ying Zhu, Ljiljana Paša-Tolić, Sindy K. Y. Tang

**Affiliations:** Department of Mechanical Engineering, Stanford University, Stanford, California, USA; Environmental Molecular Sciences Laboratory, Pacific Northwest National Laboratory, Richland, Washington, USA

**Keywords:** Tissue microdissection, Two-photon polymerization, Hierarchical blade arrays, Spatial proteomics, Mass spectrometry

## Abstract

To study tissue heterogeneity at the sub-millimeter scale, laser capture microdissection (LCM) has been the leading technology for isolating regions of interest (ROI) for downstream molecular profiling. As the ROI approaches cellular dimensions (∼10 μm), laser-induced photothermal damage and challenges in capturing microtissues in conventional LCM can compromise protein preservation and quantitative fidelity. This work introduces multi-tiered μDicers, fabricated by two-photon polymerization, to mechanically dissect tissue slices into uniform microtissues down to 10 μm. The hierarchical blade architecture limits instantaneous blade-tissue engagement and lowers the cutting force relative to single-tier designs. For benchmarking, proteomic analysis is performed on ethanol-fixed human squamous cell carcinoma microtissues generated by μDicers and by LCM. Under identical Nanodroplet Processing in One pot for Trace Samples (nanoPOTS) and liquid chromatography-mass spectrometry (LC-MS) conditions, μDicers yield more peptides and proteins than LCM, with the largest gains at 10-20 μm spatial resolution. Confocal imaging shows catapult-associated cavities in LCM-generated microtissues. This material loss, along with membrane-limited protein extraction, likely reduces protein coverage. In contrast, multi-tiered μDicers enables reproducible microdissection down to 10 μm while maintaining high protein coverage. With spatial registration of microtissues under development, μDicers have potential to complement LCM for next-generation spatial proteomic workflows.

## 1. INTRODUCTION

Biological tissues are highly heterogeneous at the cellular level. Among strategies for analyzing this heterogeneity, the physical isolation of regions of interest (ROI) prior to downstream analysis underpins key advances in histopathology and spatial proteomics.^1–5^ Laser capture microdissection (LCM) remains the most prevalent method for physical dissection of tissues at the sub-100 μm scale.^6^ In standard LCM workflows, tissue slices are first mounted on a UV-absorbing polymer membrane (e.g., polyethylene naphthalate (PEN)). The laser cuts through the membrane and the tissue, which are then collected by laser-pressure catapulting or gravity-drop.^7–9^ However, polymer membranes introduce well-documented challenges for downstream proteomics. Tissue adheres strongly to the hydrophobic membrane, hindering buffer penetration and protein extraction and thereby reducing protein coverage.^10^ Although LCM is designed to minimize tissue damage by focusing the UV laser on the membrane, direct laser interaction inevitably causes visible photodamage along the cutting path (on the order of micrometers in width), resulting in both the physical loss of tissue material within the cut path and potential molecular alterations to the adjacent regions. As the target microtissue approaches cellular scale (∼10 μm), the absolute protein mass per microtissue decreases sharply while the surface-to-volume ratio increases. Consequently, even modest losses to membranes or surfaces, together with laser-induced photothermal damage and membrane-limited extraction, increasingly compromise protein preservation and thus the ability to confidently identify and accurately quantify proteomes at micrometer scales.^11^

Mechanical dissection offers a potential route to overcome some of the challenges in LCM by avoiding UV exposure and polymer membranes. For example, blade arrays have been used to dissect tissues for proteome-scale tissue mapping.^5,12^ While these studies highlighted the potential for high throughput dissection, the spatial resolution was limited to ≥400 μm.^5,12^ At ≥400 μm, each microtissue averages the proteomic signal from many cells, which blurs microenvironmental heterogeneity and reduces sensitivity to subtle, spatially localized differences. Previously, we developed a μDicer, comprising a hollow array of microblades fabricated in silicon, to mechanically dissect tissue slices into uniform 200 μm-microtissues in a single step.^13,14^ Although the blade geometry could be tuned by silicon etching parameters and the design of the etch mask, the achievable profiles were constrained by the isotropic and anisotropic etching processes. As a result, precise control of blade tip radius below a few micrometers was challenging, and further reduction of blade tip spacing below the hundred-micrometer scale was difficult to achieve.

In this work, we advance the μDicer to enable mechanical dissection at cellular scales (i.e., 10 μm) and demonstrate its utility for proteomic analysis. Using two-photon polymerization (2PP) 3D printing, we fabricate multi-tiered, hierarchical μDicers with blade tip spacing from 10 μm to 110 μm. Previously, we showed the feasibility of 2PP to print single blades with tip radius as small as ∼100 nm to cut paraffin wax, a common tissue-embedding medium.^15^ Cutting energy was found to scale approximately linearly with total blade length. Generating smaller microtissues requires reduced blade tip spacing, which increases total blade length per unit area and consequently the required cutting energy. For example, a 10-μm spaced array has ∼10x the blade length and the required cutting energy is ∼10x more than that of a 100-μm spaced array. To overcome this challenge, we design a multi-tiered blade array that reduces the cutting energy by limiting the total blade length engaged with the tissue at a given time. As the tissue is pushed through the multi-tiered blade array, each tier progressively cuts the microtissues into smaller pieces until the desired size is achieved (Figures 1 and S1). By distributing the cut across multiple tiers, this hierarchical design alleviates the force-length scaling limitations of parallel, high-density blade arrays to enable controlled dissection at cellular scales. As a proof of concept, we apply the multi-tiered μDicers to dissect 10 μm-thick hematoxylin and eosin (H&E) stained, ethanol-fixed tissue sections from human squamous cell carcinoma (SCC). We evaluate the cutting forces of the multi-tiered μDicers against single-tiered μDicers and demonstrate the generation of microtissues with lateral dimensions of 10, 20, 35, and 110 μm respectively. The dimensions of these microtissues correspond to the spacings between the blade tips in the μDicers used to generate them. The resulting microtissues are collected in Nanodroplet Processing in One pot for Trace Samples (nanoPOTS) chips and analyzed by Liquid Chromatography-Mass Spectrometry (LC-MS) following published protocols.^1^ Compared with conventional LCM using laser-pressure catapulting, microtissues generated by μDicers yield higher proteome coverage with greatest improvement observed at 10 −20 μm spatial resolution.

**Figure 1.**
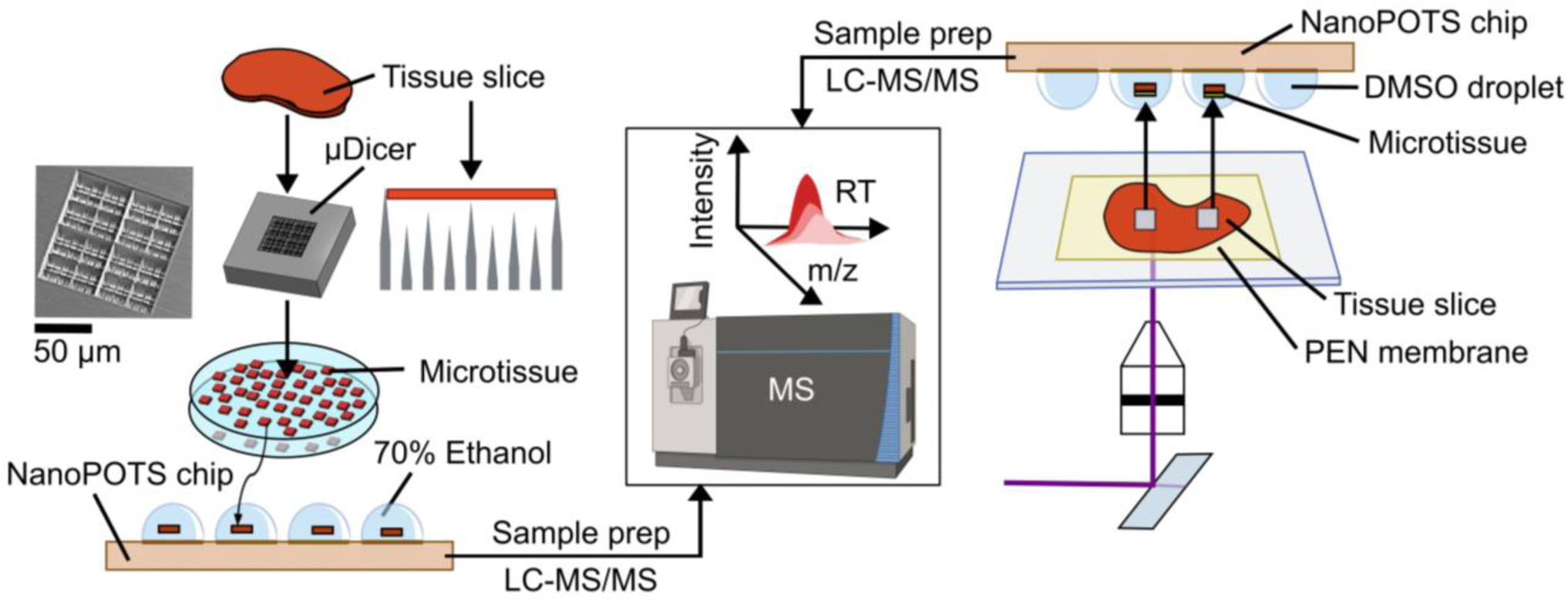
Microtissue generation and transfer to nanoPOTS chip for proteomic analysis. Left: Illustration of tissue microdissection in a multi-tiered μDicer and transfer to nanoPOTS wells prefilled with 70% ethanol. Right: Illustration of tissue microdissection and catapulting into nanoPOTS chip preloaded with DMSO droplets using LCM. Middle: Subsequent LC-MS proteomics and data analysis using TIFF (Transferring Identification based on FAIMS Filtering) approach.

## 2. RESULTS AND DISCUSSION

### 2.1 Fabrication and Characterization of Multi-tiered μDicers

The multi-tiered μDicers were designed with a vertical offset of 20 μm between the blade tips of consecutive tiers, approximately twice the thickness of the SCC tissue slices used. This design allowed the tissue to be cut by the top tier of blades before engaging the next tier. Scanning Electron Microscopy (SEM) images verified the fidelity of the μDicers printed by the Nanoscribe (Figures 2 and S2-S5), including the targeted blade tip diameter of 1 μm. We chose to use a blade tip diameter of 1 μm because it could be printed reliably with a 25x objective on the Nanoscribe and was sufficiently sharp to cut through H&E stained, ethanol-fixed SCC tissue slices and other types of tissue and materials tested (Figure S6).

**Figure 2.**
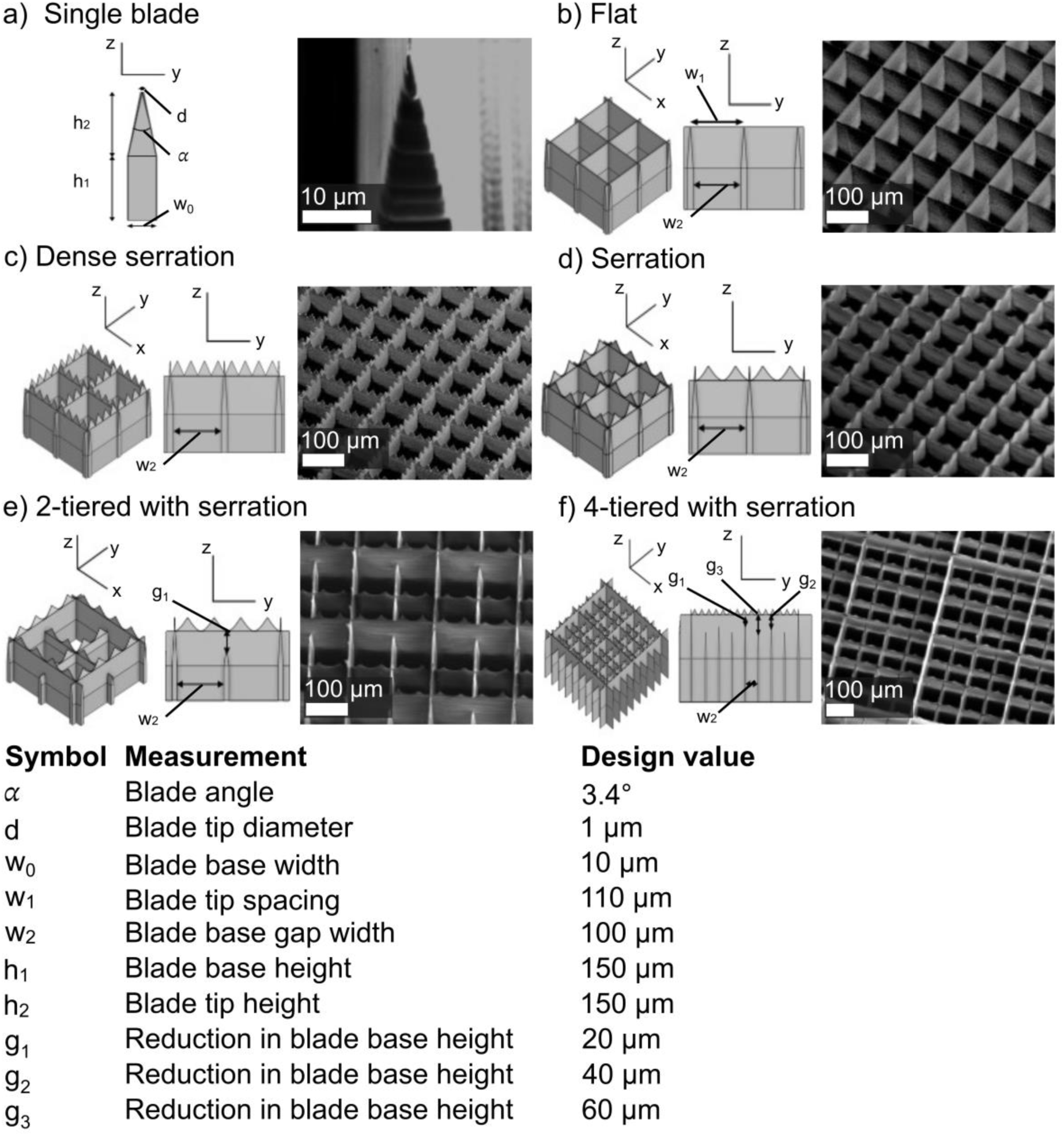
Computer-aided designs (CADs) (left panels) and SEM images (right panels) of μDicers printed with two-photon polymerization. a) Single blade. b-f) μDicers with a blade tip spacing of 110 μm (x–y direction). b) Flat: μDicer with a single tier of flat blades. c) Dense serration: μDicer with a single tier of dense serrations, consisting of 4 serrations per blade unit (110 μm long). d) Serration: μDicer with a single tier of serrations, consisting of 2 serrations per blade unit (110 μm long). e) 2-tiered μDicer with serration: the second tier was lowered by g_1_ = 20 μm. f) 4-tiered μDicer with serration: the tiers were lowered by g_1_ = 20, g_2_ = 40, and g_3_ = 60 μm respectively. The bottom table lists the blade dimensions.

To evaluate the effectiveness of the hierarchical blade architecture, we obtained force-displacement curves of the μDicers indented into H&E stained, ethanol-fixed SCC tissue slices mounted on wax-coated glass slides (Figure 3a). The maximum indentation depth was 100 μm to ensure complete cutting of 10 μm-thick tissue sections and a portion of the wax. After indentation, visual inspection of the μDicer revealed that the tissue/wax was cut and the wax became partially detached from the substrate and transferred to the μDicer (Figure 3b). Figures 3c and S7 show representative force-displacement curves of μDicers with different blade geometries. We used the maximum force required to indent 100 μm of tissue/wax as a practical metric for accessing and comparing various blade geometries. Since μDicers designed to generate different microtissue sizes had varying blade array areas in this set of experiments (refer to Section 4 – Experimental Methods for details), comparisons were limited to μDicers designed for the same microtissue size. While the resolution of the force measurement system we used was insufficient to delineate the detailed features of the cutting process (e.g., saw-tooth patterns indicative of stick-slip motion during cutting),^16,17^ we expect that the creation of new surfaces, plastic deformation, and friction between the blades and the tissue/wax all contributed to the measured force.^15^

**Figure 3.**
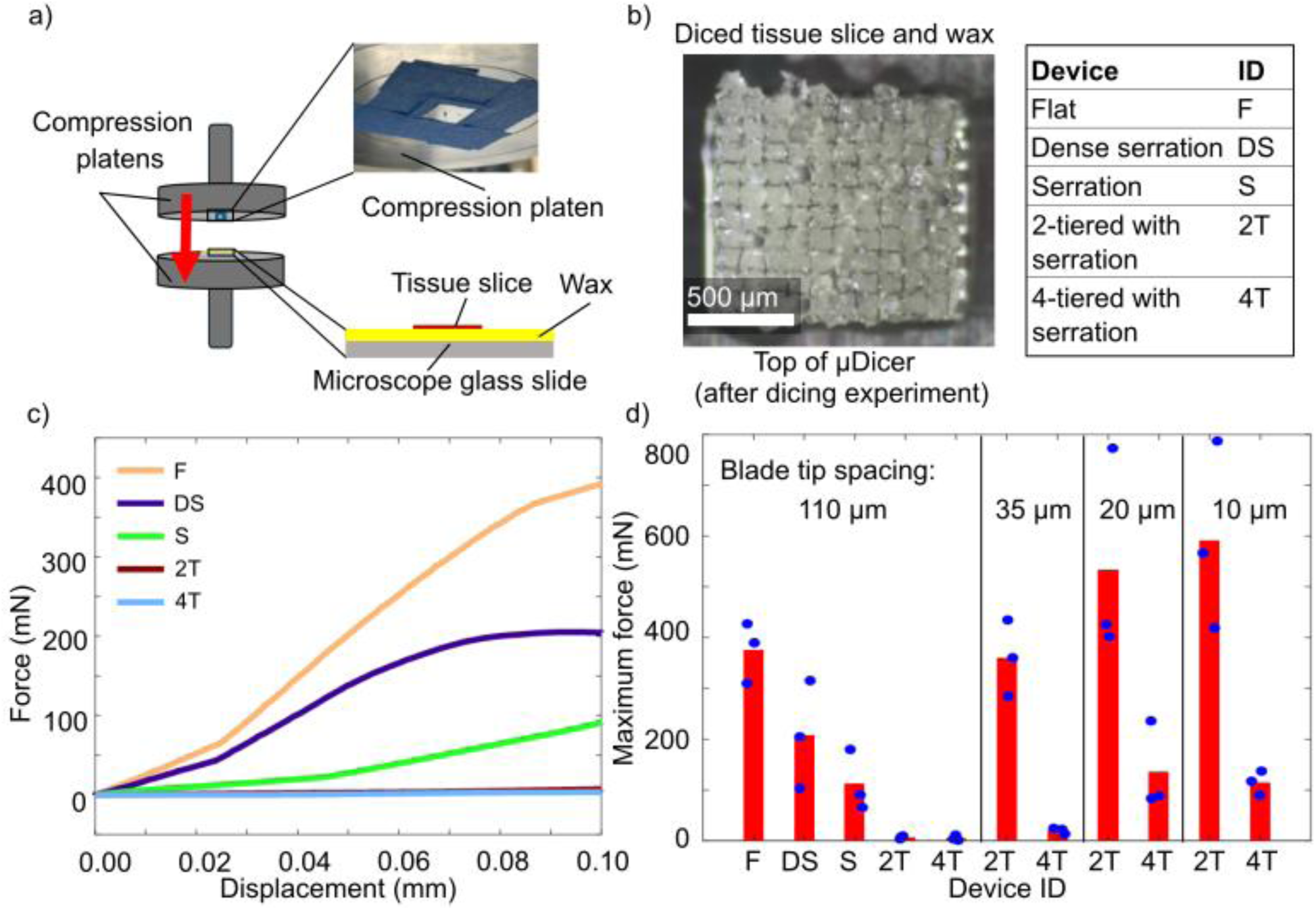
Force-displacement characterization of μDicers. The substrate was a 10 μm-thick hematoxylin and eosin (H&E) stained, ethanol-fixed squamous cell carcinoma (SCC) tissue slice. a) Experimental setup with μDicer and SCC tissue slices on wax-coated glass slide mounted on the top platen and the bottom platen of the Instron, respectively. b) Post-dicing image of the top of the μDicer (blade tips facing up) showing cut wax pieces. Cut microtissues were not visible because they were inside the μDicer underneath the wax. c) Force-displacement curves for μDicers with different blade geometries. The μDicers here had a blade tip spacing of 110 μm and a blade array area of 1 mm x 1 mm. d) Graph showing the maximum cutting force at a displacement of 0.1 mm. Each data point was measured at a different location on the same SCC tissue slice (derived from soft tissue). For 2-tiered and 4-tiered μDicers with 10 μm blade tip spacing, we used SCC tissue slice derived from lymph node to collect the data. Each blue marker represents one measurement. The height of the red bars represents the mean for each condition.

Among single-tiered μDicers with a blade tip spacing of 110 μm, serrated blades required the lowest maximum force (∼30% of that needed for flat blades) (Figure 3c). This outcome was expected due to stress concentration at the tips of the serrated blades.^18,19^ In addition, our system was likely dominated by friction. The reduced contact area of the serrated blades with the tissue/wax resulted in a lower maximum force compared with densely serrated blades. Among serrated μDicers, increasing the number of tiers of blades further reduced the maximum cutting force with the effect being most pronounced in μDicers with blade tip spacing below 35 μm. In μDicers with 20 μm and 10 μm blade tip spacings, the mean maximum force for the 4-tiered design was only 26% and 19% respectively, of those observed for the 2-tiered design. This force reduction aligns with the hypothesis that the multi-tier architecture reduces the blade area interacting with the tissue slice. Although the cutting sequence could not be observed directly, the 20 μm inter-tier offset (approximately twice the tissue slice thickness) facilitated sequential cutting, where the top tier initially cut the tissue slice to free the microtissue edges before the subsequent tiers engaged with the tissue. This sequential engagement reduced tissue stress and enabled more efficient microtissue generation than the single-tiered design, which must deform and cut the tissue simultaneously into small microtissues.

To test reusability of the 3D printed multi-tiered μDicers, we used a single flat blade and performed two indentations on different regions of the same tissue slice mounted on a wax-coated slide. After the first indentation, the blade tip dulled and the maximum cutting force more than doubled (Figure S8). Accordingly, we used our μDicers only once in all subsequent experiments. This practice also prevented cross-contamination between devices.

### 2.2 Characterization of Microtissues Generated by the μDicers

Because the 4-tiered μDicers with serrated blades achieved the lowest maximum force in indentation experiments, they were used in all subsequent experiments. To evaluate the size distribution of the microtissues generated by the μDicers, we transferred a circular section of H&E stained, ethanol-fixed SCC tissue (2 mm in diameter and 10 μm in thickness) to the top of the μDicer and pushed the tissue through the μDicer using a biopsy punch plunger modified with a 2 mm-thick silicone tip (Figure 4a,b; refer to Section 4 – Experimental Methods for details). Figure 4c-e show images of the SCC microtissues generated by the 4-tiered μDicers with blade tip spacings of 110 μm, 35 μm, 20 μm, and 10 μm, respectively. Cell nuclei are visible in individual microtissues (as indicated by red arrowheads). Irregularity in the shape of some microtissues may arise from the heterogeneity in local tissue stiffness, slight variations in the slice thickness, and/or folding of the microtissue edges. These irregularities were more prominent in 110 μm microtissues due to their small thickness-to-width ratio. As measured by confocal microscopy (refer to Section 4 – Experimental Methods for details), the volumes of the microtissues generated by all tested μDicers matched the expected volumes (Figure 4f).

**Figure 4.**
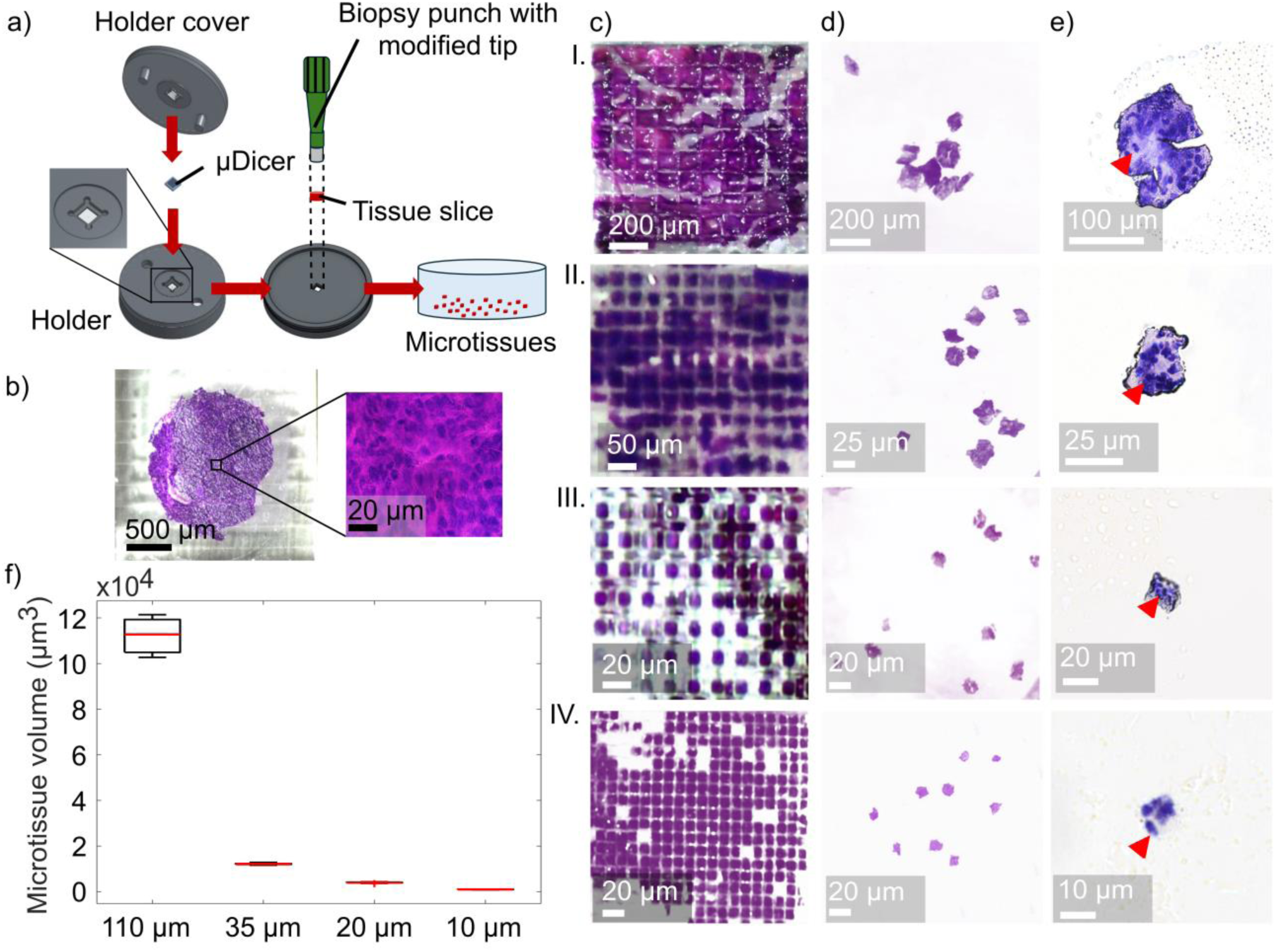
Generation of SCC microtissues using 4-tiered μDicers. a) Setup to push the tissue slice through the μDicer using a biopsy punch with a modified tip. b) Photograph of a H&E stained, ethanol-fixed SCC slice on top of a μDicer. Cell nuclei are visible in the magnified image (right panel). c) Diced microtissues before their release from the μDicers: bottom views (I–III) of μDicers with 110, 35, and 20 μm blade tip spacings (with through-holes); top view (IV) of μDicer with 10 μm blade tip spacing (without through-holes). d) Microtissues released in 70% ethanol. e) Magnified images of SCC microtissues generated by μDicers with blade tip spacing of 110 μm, 35 μm, 20 μm, and 10 μm, respectively (I-IV). Red arrows indicate visible nuclei. f) Box plot of microtissue volumes (n = 10). Red horizontal lines indicate medians. Boxes indicate the interquartile range (IQR, 25th to 75th percentile), and whiskers extend to the most extreme data points that are not considered outliers.

We evaluated whether it was possible to generate microtissues of similar sizes using an alternative mechanical dissection method, specifically a commercial tissue chopper (McIlwain) mounted with a stainless-steel razor blade. We observed excessive heat generation due to the reuse of the blade to apply a normal force on the tissue slice. This heat likely led to inaccurate blade displacement. A significant portion of the tissue slice remained uncut, while the microtissues that were produced exhibited poor adhesion to the glass slide and were scattered during the cutting process. Hence, only a limited number of microtissues with a lateral dimension of 110 μm were collected and none with a lateral dimension of 25 μm or smaller were obtained (Figure S9).

### 2.3 Proteomic Performance of LCM vs. μDicer Microdissection

Finally, to compare proteome coverage attainable with LCM vs. μDicers, we performed quantitative proteomic analyses of microtissues generated by each method at different spatial resolutions: 10 μm, 20 μm, and 110 μm. All samples were collected into nanoPOTS chips and processed using identical on-chip digestion protocol and LC-MS conditions (see Section 4 – Experimental Methods for details). Transferring Identification based on FAIMS Filtering (TIFF)-based MS data acquisition and MSFragger were used to process LC-MS data.^20^ The TIFF approach integrates ion mobility into LC-MS workflows to improve peptide identification in low-input samples, where limited ion abundance and sampling stochasticity can reduce tandem MS (MS2) coverage. To address these challenges, bulk library samples were run concurrently to enable confident MS2-based peptide identification. Peptide features in microtissues were then identified using a spectral library and an MS1-based match-between-runs (MBR) strategy.

Across all microtissue sizes, μDicer yielded more unique proteins and peptides than LCM (*p* < 0.001, indicating statistically significant differences), with the largest gains at 10 μm and 20 μm (Figures 5a,b and S10). Protein intensities scaled with microtissue size for both methods and were consistently higher for μDicer at matched sizes (median log2 intensities for μDicer vs. LCM: 17.4 vs. 15.1 at 10 μm; 18.6 vs. 16.1 at 20 μm; 22.5 vs. 21.4 at 110 μm; Figure 5c). Differences in the proteome coverage were even more pronounced when considering identifications based on MS2 instead of MS1-based MBR approach commonly used for low-input proteomics (Figure S10). The fraction of fully tryptic peptides and methionine-oxidized peptides were similar for 110 μm microtissues generated by LCM and μDicer (Figure S11). We anticipated similar fractions of tryptic and methionine-oxidized peptides in smaller microtissues based on previous studies comparing LCM and deep-UV ablation.^11^ These results suggest that the improved proteome coverage achieved by the μDicer is unlikely to stem from variations in digestion efficiency or the presence of oxidation artifacts.

**Figure 5.**
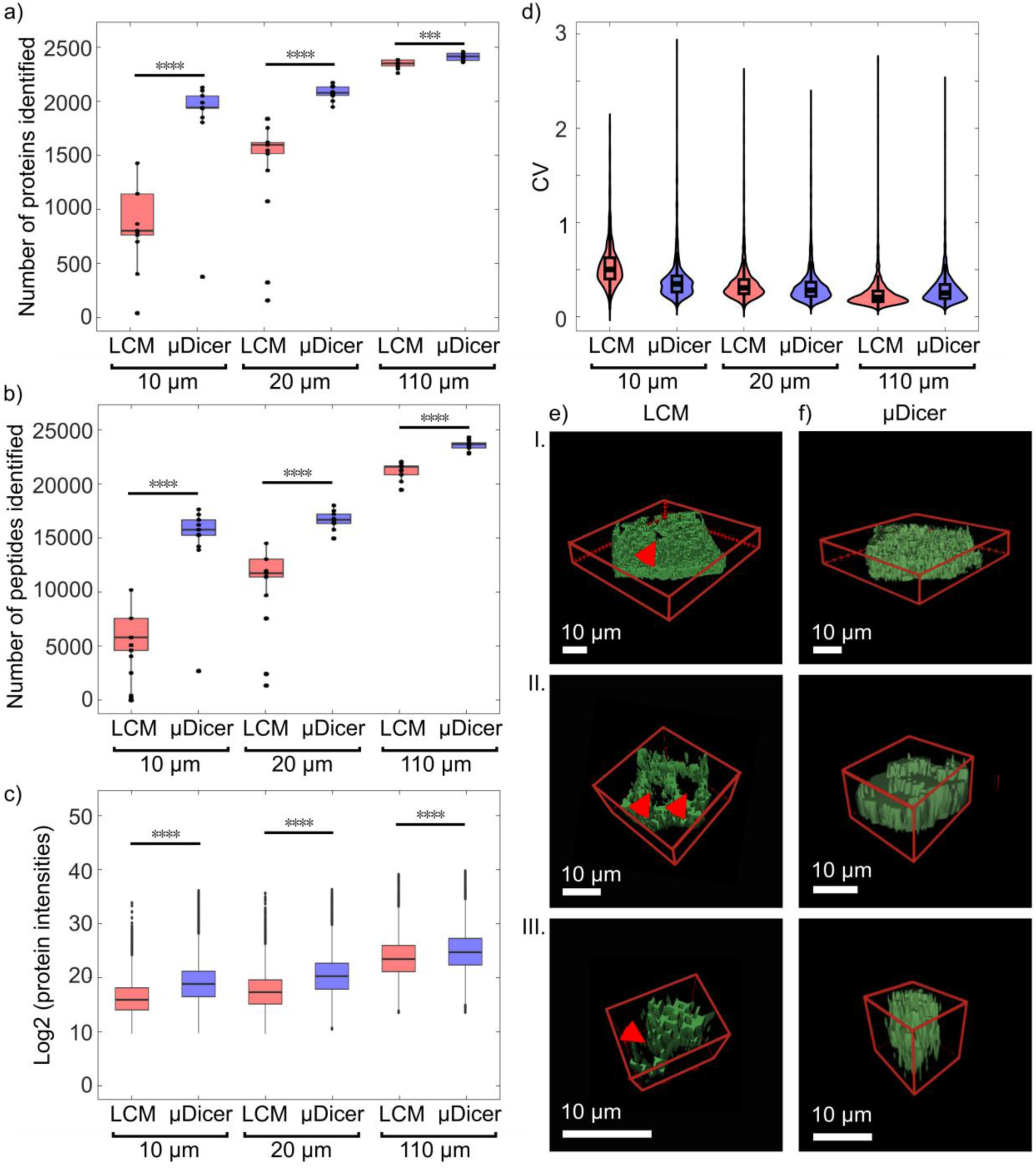
a) Number of proteins and b) peptides quantified from SCC microtissues of different sizes, generated using LCM and μDicers. c) Log2 protein intensities of microtissues generated by LCM and μDicers. In a)-c), Wilcoxon rank-sum test was used for the comparisons between LCM vs. μDicers: *** *p* ≤ 0.001; **** *p* ≤ 0.0001. d) Violin plots indicating the distributions of coefficients of variations (CVs) of protein intensities across microtissues of various sizes. 3D rendering of confocal images of SCC microtissues generated by e) LCM and f) μDicers at spatial resolutions of 110 μm, 20 μm, and 10 μm, respectively (I-III). The LCM-generated microtissues exhibit deformation and material loss, particularly in 20 and 10 μm microtissues. Red arrowheads indicate through-holes or cavities.

To evaluate the quantitative performance attainable by LCM vs. μDicer across spatial resolutions, we performed pairwise Pearson correlations of protein intensities (Figure S12). As expected, larger microtissues exhibited higher correlations in both LCM (r=0.95 at 110 μm) and μDicer (r=0.94 at 110 μm) proteomic analyses. Differences emerged at smaller sizes: at 10 and 20 μm, while LCM replicates showed broader variability (r=0.57-0.69), μDicer replicates remained tightly correlated (r=0.83-0.92). The lower correlation coefficients for the smallest microtissues regardless of the method may be attributed to the higher tissue heterogeneities near cellular scale. After excluding proteins with >50% missing data per group, we also compared median coefficients of variations (CVs) of protein intensities across replicates for two methods (Figure 5d). Median CVs for LCM-generated microtissues were 0.180 (110 μm), 0.291 (20 μm), and 0.505 (10 μm), while μDicer median CVs were 0.232 (110 μm), 0.264 (20 μm), and 0.337 (10 μm). Consistent with correlation analysis, μDicer yielded lower CVs at 10 and 20 μm, indicating improved quantitative reproducibility, reduced technical variability, and increased detection reliability of subtle proteomic differences at higher spatial resolution (i.e., smaller microtissues) compared with LCM.

Among microtissues generated by the μDicer, proteins identified at 110, 20, and 10 μm showed a large three-way overlap, indicating consistent proteome capture across these microtissue sizes (Figure S13). When we performed the same “across-sizes” comparison within each dissection method, the common set of proteins shared by 110, 20, and 10 μm was larger for μDicer than for LCM, suggesting that μDicer maintained proteome coverage with reduced microtissue sizes (Figure S13a,b). At 10 μm, microtissues generated by LCM and μDicer shared a substantial fraction of proteins, with additional proteins unique to μDicer (Figure S13c).

To investigate the potential cause of the discrepancy between LCM and μDicer proteomic outcomes, we compared confocal z-stacks of microtissues from both methods (Figure 5e,f). At matched target microtissue sizes, LCM-generated microtissues exhibited markedly smaller volumes than those generated by μDicer. In representative LCM-generated microtissues, the measured volumes were 51% of the expected volumes at 110 μm, 48% at 20 μm, and 25% at 10 μm. However, in representative μDicer-generated microtissues, the measured volumes were much closer to the expected volumes (i.e., 93% at 110 μm, 99% at 20 μm, and 92% at 10 μm) (Figures 5e,f and S14, Table S1). The volume losses in LCM-generated microtissues coincided with cavities in both the microtissues and the underlying PEN membranes, consistent with catapult-induced material loss during tissue collection. Combined with the known reduction in protein extraction efficiency on PEN membrane,^10^ the diminished volume and membrane effects provide a mechanistic explanation for the lower protein coverage observed with LCM. To our knowledge, this discrepancy between the expected and observed volumes of LCM-generated microtissues has not been previously reported.

## 3. CONCLUSION

As spatial proteomics advances towards cellular scales, preserving protein content during microdissection becomes essential for achieving deeper proteome coverage, improved quantitative accuracy, and reproducibility. LCM has long been the predominant technology for sub-100 μm tissue dissections due to its high precision and minimal invasiveness. However, its efficacy below 20 μm is often limited by factors such as laser spot size, localized thermal and photochemical effects, polymer membrane chemical properties, and tissue damage during catapulting. In this study, catapult-associated effects such as cavities observed in the microtissue and PEN membrane coincided with noticeable reduction in proteome coverage. Alternative LCM modalities that bypass the use of PEN membranes and physical catapulting, including deep-UV laser ablation^11^ and gravity drop may help address some of these issues, but their ability to achieve good proteomic performance at ≤ 20 μm resolution needs to be carefully evaluated.

To our knowledge, this work is the first demonstration of uniform, reproducible mechanical microdissection of human tissue slices with lateral dimensions down to 10 μm. Unlike tissue choppers, the μDicer generates all microtissues in a single, parallel pass and avoids repeated impact that heats the sample and rapidly blunts the blade. The multi-tiered μDicer architecture employed herein was key to enabling dissection at 10 μm by limiting the blade length engaged with tissue at any instant, thereby reducing tissue deformation and the required cutting force. Mechanical dissection eliminates UV exposure and polymer membranes, thus reducing sample loss, photodamage, and membrane-related inefficiencies that can compromise protein recovery. The performance superiority of the μDicer over LCM became more evident as microtissue sizes approached cellular scales (10 −20 μm).

The μDicer currently produces uniform microtissues but does not yet support targeting regions of interest. Ongoing developments aim to enable spatial registration, integrate multiplexed workflows, and improve throughput with optimized LC methods and automated processing, paving the way for high-throughput, high-resolution 3D organ mapping through spatial proteomics.

## 4. EXPERIMENTAL METHODS

### Squamous cell carcinoma tissue slice preparation

We collected de-identified fresh frozen human squamous cell carcinoma (SCC) from the parotid gland, soft tissue, lymph node, and skin from the Stanford Tissue Bank and stored the samples at −78 °C until use. The tissues obtained from the Stanford Tissue Bank were excess specimens not needed for clinical use, collected from appropriately consented patients who underwent standard-of-care medical procedures unrelated to the study. The Stanford Institutional Review Board (IRB) determined that this study does not meet the definition of clinical investigation under 21 CFR 50.3(c) and that this study does not involve human subjects as defined in 45 CFR 46.102(e) and 21 CFR 50.3(g).

We used a cryostat (LEICA CM3050 S) to section the SCC tissue samples into 10 μm slices without any embedding media. We collected the tissue slices on top of polydimethylsiloxane (PDMS)-coated glass slides within 30 seconds of sectioning inside the cryochamber of Cryostat (−20 °C). The SCC slices on top of the PDMS-coated glass slides were then stored on dry ice (−78 °C) for up to 30 minutes before further tissue processing (i.e., ethanol fixation and H&E staining). Prior to processing, the slides were left at room temperature (∼22 °C) for one minute. The SCC slices remained sufficiently adhered to the PDMS-coated glass slides for subsequent manipulations.

We fixed the SCC tissue slices using 50% ethanol (30 seconds), 70% ethanol (30 seconds), 90% ethanol (30 seconds), and 95% ethanol (30 seconds) respectively. All ethanol solutions were pre-chilled at 4 °C before fixation which occurred at room temperature. We stained the ethanol-fixed SCC slices using a conventional hematoxylin and eosin (H&E) staining protocol which was performed at room temperature. Briefly, we applied sufficient hematoxylin to cover the tissue slices on PDMS-coated glass slides in a Petri dish for 3 minutes. Then we rinsed the tissue slices in two changes of deionized (DI) water to remove excess stain. Next, we applied sufficient bluing reagent to cover the tissue slices for 15 seconds, followed by rinsing in two changes of DI water. Next, we dipped the tissue slices in 99% ethanol. Finally, we applied sufficient eosin Y solution – modified alcohol to cover the tissue slices for 2 minutes, followed by rinsing and dehydration using 99% ethanol. The H&E stained, ethanol-fixed SCC tissue slices were then dried using compressed air, followed by vacuum sealing and storage at −78 °C to keep the tissue dehydrated until use. We measured the Young’s modulus of the H&E stained, ethanol-fixed SCC tissue slices to be on the order of 100 kPa (Figure S15).

### Fabrication of μDicers for the generation of microtissues

We printed our μDicers with the Nanoscribe Photonic Professional GT or Quantum X shape using a 25x objective in IP-S photopolymer resin on an indium tin oxide (ITO) coated glass substrate (Figure S2). The blade tip diameter was designed to be 1 μm. We varied the blade tip spacing from 10 μm to 110 μm. All blades had a base width of 10 μm. The thickness (in the z direction indicated in Figure 2 and S3-S5) of the μDicer device was 300 μm. The μDicers with blade tip spacings ≥ 20 μm had through-holes (see details in Figures S3,4). Due to the finite width of the blade base (10 μm), the μDicer with blade tip spacing = 10 μm had no through-holes (see details in Figure S5). The μDicers with 10, 20, and 35 μm blade tip spacings were designed to simultaneously generate a maximum of 2500 microtissues in a 50 x 50 blade array, while the μDicer with a 110 μm blade tip spacing was designed to generate a maximum of 100 microtissues in a 10 × 10 blade array. μDicers were developed using SU-8 developer (15 minutes) and cleaned using isopropanol (5 minutes). For the μDicers with 10 μm blade tip spacing only, we used a prolonged developing and cleaning protocol (i.e., 30 minutes of immersion in SU-8 developer, 30 minutes of immersion in isopropanol, followed by 30 minutes of immersion in isopropanol in an ultrasonic (∼25 kHz) bath) to remove any residues of the resin at the bottom of the μDicer. After development and cleaning, all devices were air dried for 5 minutes. Next, we detached the μDicer from the ITO coated glass substrate manually using a tweezer. To reduce friction during dicing, we coated the surface of the μDicers with Aquapel™ (30 minutes) and FC40 (30 minutes). The device was then washed in DI water (5 minutes), followed by rinsing in 70% ethanol and air drying for generating the microtissues.

### Characterization of the μDicers

To evaluate the fidelity of the prints, we used a scanning electron microscope (SEM) (Thermo Fisher Scientific Apreo S) to image the μDicers, which were sputter-coated (Kurt J. Lesker) with a 10 nm layer of gold prior to imaging.

To evaluate the force-displacement characteristics of the μDicers, we used Instron 5565 and a 100 N load cell. For these measurements, we used tweezers to place the SCC tissue slices (10 μm-thick) on glass slides coated with ∼1 mm of paraffin wax (Fisherbrand^TM^ Histoplast Paraffin Wax, catalog # 22-900-700). Briefly, we melted paraffin wax in a glass beaker at 70 °C. We then poured a small amount of the melted wax on top of the glass slides. We scraped the melted wax gently using the blunt side of a razor blade to distribute the melted wax on the glass slides. Finally, we let the wax cool at room temperature until it completely solidified (∼15 minutes). After generating 10 μm SCC tissue slices using Cryostat, we placed the tissue slices on top of wax-coated glass slides within 30 seconds inside the cryochamber of Cryostat (−20 °C). We stored the SCC slices on top of wax-coated glass slides on dry ice (−78 °C) for up to 30 minutes and performed ethanol fixation and H&E staining using the same protocol as described before. Finally, the force-displacement characteristics were performed on the H&E stained, ethanol-fixed SCC slices on top of wax-coated glass slides.

For the Instron experiments only, we attached the μDicer on ITO coated glass on the top compressor platen and SCC tissue slice on wax coated glass slide on the bottom platen using blue masking tape. We used a constant displacement rate of 0.20 mm/minute and measured the force for a 100 μm-displacement. For these experiments only, the μDicer were not coated with Aquapel™ or FC40. The blade tips in the μDicer were designed to be 300 μm taller than the rim of the device to facilitate the indentation of the μDicer blades into the substrate. To reduce print time in the Nanoscribe, the μDicer blade array area was made smaller than the array area used to generate microtissues. Specifically, the blade array areas for μDicers with blade tip spacings of 10 μm, 20 μm, and 35 μm were 80 μm x 80 μm, 160 μm x 160 μm, and 200 μm x 200 μm, respectively. These designs still allow us to evaluate the effect of blade geometries and the number of blade tiers for each given blade tip spacing (to generate a given microtissue size), while reducing print time in the Nanoscribe.

### Generation of microtissues using the μDicers

Because the μDicer was 300 μm thick only, to support and stabilize the μDicer during dicing experiments, we sandwiched the μDicer between a holder and a holder cover (Figure 3a), which were 3D-printed using Elegoo Saturn 2 in Standard Photopolymer Resin (Black). To transfer a tissue slice onto the μDicer, we used a biopsy punch to extract a 2 mm-circular section from the H&E stained, ethanol-fixed SCC tissue slice on the PDMS-coated glass slide. We then transferred the tissue slice on top of a μDicer. To dice and push the tissue through the μDicer without damaging the blades, we pressed on the tissue slice using a 2 mm-diameter biopsy punch plunger with a modified tip comprising a 2 mm-thick silicone (Dragon Skin™ 10, Young’s Modulus ∼90 kPa). To facilitate the recovery of microtissues, we immersed the μDicer with any adhered microtissues in 70% ethanol in a Petri dish inside an ultrasonic bath (∼25 kHz) for 2 minutes (Figure S16).

### Characterization of microtissue sizes

We used brightfield microscopy (Zeiss Stemi 508 (5x objective) or Keyence BZ-X800 (20x objective)) or confocal microscopy (Zeiss LSM-780 confocal microscope (40x objective for microtissues generated using μDicer with blade tip spacings ≥ 20 μm and 63x objective for blade tip spacings = 10 μm)) to characterize the size distribution of the microtissues. We analyzed the images using Fiji (Image J). For phantom tissues i.e., agar (2%) and wax as well as fixed tissues i.e., beef liver and human prostate tissue grown in a mouse (LTL 610), we determined microtissue widths by considering the narrowest dimensions passing through the center of the microtissues (Figure S6). For H&E stained, ethanol-fixed SCC microtissues, we determined microtissue volumes from confocal z-stacks: images were segmented by intensity thresholding, objects identified by connected-component labeling, and volumes calculated as voxel count × voxel volume. Because the tissues were autofluorescent, we were able to image them in the Alexa Fluor 488 channel. The PEN membranes had a different autofluorescence spectra and were clearly differentiated from the tissue in the DAPI channel.

### Fabrication of nanoPOTS chips

NanoPOTS chips containing an array of 4 x 12 nanowells were used, where each nanowell had a diameter of 1.2 mm and a center-to-center spacing of 4.5 mm. Glass slides, precoated with chromium and photoresist (25 mm x 75 mm, Telic company) were used for chips fabrication using a previously published protocol.^1^ Following wet etching, the glass surfaces were treated with 2% (v/v) heptadecafluoro-1,1,2,2-tetrahydrodecyldimethylchlorosilane in 2,2,4-trimethylpentane. The remaining chromium layer was then removed to reveal an array of hydrophilic surfaces for tissue collection and proteomic sample preparation.

### Transferring microtissues onto nanoPOTS

For mass spectrometry analysis, microtissues generated by the μDicers were collected in a Petri dish containing 70% ethanol and transferred at room temperature using a micropipette. Each microtissue was transferred into an individual well on the nanoPOTS chip along with 250 nL of 70% ethanol. After transferring the microtissues into nanoPOTS chip, we left the chip in a laminar flow hood for 15 minutes to air-dry the H&E stained, ethanol-fixed microtissues completely. The nanoPOTS chip, covered with a custom plastic lid fitting the chip’s edges with a central hollow, was wrapped with parafilm followed by aluminum foil, and subsequently vacuum-sealed in a plastic bag. We used overnight shipping to ship the nanoPOTS chip to Pacific Northwest National Laboratory (PNNL) in 1 kg of dry ice in a Styrofoam box. Upon receipt, the samples were preserved at −78 °C until mass spectrometry analysis.

### Laser capture microdissection: collection of microtissues onto nanoPOTS

To benchmark the μDicer method, a consecutive tissue slice, mounted on a PEN membrane slide, was processed via standard LCM-nanoPOTS workflow to serve as a direct comparison in the mass spectrometry analysis. Briefly, tissue slices with 10 μm thickness were mounted onto PEN membrane coated slides (Carl Zeiss). 200 nL of dimethyl sulfoxide (DMSO) was loaded onto each nanoPOTS well. A PALM Microbeam IV LCM system (Carl Zeiss Microimaging) was used for LCM and collection of tissues. Square regions of 10 × 10, 20 x 20, and 110 x 110 μm^2^ were generated from the consecutive slice and collected directly into DMSO droplets deposited on nanoPOTS wells. An energy level of 35 was used to dissect the tissue pixels and collected onto the NanoPOTS using catapult energy of 15 and focus energy of delta 4. After collection, DMSO was evaporated by heating the chip to 70 °C for 20 minutes. The dried chip was inspected using a microscope to ensure successful microtissue collection. The chip was stored at −20 °C until proteomics sample preparation.

### Proteomics sample preparation using a nanoPOTS platform

μDicer and LCM generated microtissues were processed in parallel on the nanoPOTS platform for mass spectrometry analysis. A robotic liquid-handling platform, described elsewhere, was used for preparation of proteomics sample preparation.^21^ Extraction of proteins followed by reduction was performed using a lysis buffer containing 0.1 % (v/v) n-dodecyl-β D-maltoside, 1 mM tris(2-carboxyethyl)phosphine and 50 mM ammonium bicarbonate (ABC) (pH 8.0). 200 nL of lysis buffer was dispensed into each well and incubated for 1 hour at 70 °C. Next 50 nL of 10 mM of 2-iodoacetamide in 50 mM ABC buffer was added to each well and incubated in the dark for 30 minutes at room temperature. Following alkylation, the proteins were digested overnight using a 50 nL solution containing 0.5 ng Lys-C (MS grade, Promega) and 2 ng of trypsin (Promega) per well. The nanoPOTS chips were wrapped with aluminum foil and incubated in a hydration chamber during the entire protocol to ensure minimal droplet evaporation. After overnight digestion, the reaction was quenched by adding 50 nL of 5% formic acid (FA) into each well. The droplets were then evaporated in a vacuum desiccator and stored at −20 °C until further analysis.

### LC-MS/MS analysis

A home-built nanoPOTS autosampler platform was used to sequentially perform sample collection, cleanup and chromatographic separation.^22^ An in-house column (100 μm i.d., 4 cm) packed with C18 (300 Å pore size; Phenomenex) column was used as solid phase extraction column. For chromatographic separation, an in-house packed LC column (50 μm i.d., length 25 cm, packed with 1.7 μm, C18 packing material (BEH 130 Å C18 material, Waters) was used. A column heater (Analytical Sales and Services Inc.) was used to maintain a constant column temperature of 50 °C during separation. Buffer A (0.1% FA in water) was used to dissolve the dried peptides on the nanoPOTS chips and injected into the solid phase extraction column for 5 minutes using 2% acetonitrile as the loading buffer. The LC-MS system consisted of an UltiMate 3000 RSLCnano System (Thermo Fisher Scientific) coupled with an Orbitrap Lumos tribrid MS (Thermo Fisher Scientific). The peptides were then separated using gradient of buffer B (0.1% FA in acetonitrile), eluted at a flow rate of 100 nL per minute. A 30-minute linear gradient (from 8% to 22%) of buffer B followed by a 9-minute linear gradient (from 22% to 35%) of buffer B was used for separation. Eluted peptides were ionized using electrospray ionization source, operated at a voltage of 2400 V.

A discrete field asymmetric waveform ion mobility spectrometry (FAIMS pro) interface was used to enhance the sensitivity of the analysis by filtering out singly charged species.^20^ Spectral libraries were constructed by sequentially analyzing 50 ng of digested peptides from dissociated SCC tissues at three different compensation voltages (−45, −60 and −70) in three separate LC-MS runs. Fractionated peptide ions were first analyzed using orbitrap using the following parameters: mass range (m/z) 350 to 1500, resolution: 120,000, ion injection time: 120 ms, automatic gain control (AGC) target: 1E6. Precursor ions with charges ranging from +2 to +6 and intensities > 1E4 were selected for further tandem MS2 analysis. The selected ions were isolated with a mass window of 1.4 m/z and fragmented with 30% level of high energy collisional dissociation (HCD). MS2 fragment ions were then analyzed in an ion trap operated at a maximum injection time of 86 ms and AGC of 2E4. The total cycle time was set to 2 s. For analysis of the samples, three FAIMS compensation voltages (−45, −60 and −75) were used in three consecutive scans for each LC-MS run. For MS1 analysis of the fractionated peptides in the orbitrap, the following parameters were used: mass range (m/z) 350 to 1500, resolution: 120,000, max ion injection time: 500 ms, AGC target: 1E6. Peptide precursor ions with charge states ranging from +2 to +6 and intensities greater than 1E4 were selected for MS2 analysis. The following parameters were used for MS2 analysis in the ion trap: isolation window: 1.4, HCD Energy: 30% normalized level, max IT: 150 ms, AGC: 2E4.

### MS data analysis

Fragpipe (v 23.1), equipped with MSFragger search engine, Philosopher for false discovery rate (FDR) filtering and reporting, and IonQuant for label-free-match-between-runs (MBR) and quantification were used for processing raw MS spectrum files. The following parameters were used for search: full tryptic specificity up to 2 missed cleavage sites; fixed modifications: carbamidomethylation (57.0214 Da) on cysteine, and methionine oxidation (15.9949); variable modifications: protein N-terminal acetylation (42.0105 Da). Peptides were identified and matched to the reference library using retention time, *m*/*z,* and FAIMS CV. An FDR rate of 0.05 was used during peptide search. Data cleanup, statistical analysis, and visualization were performed using R Studio (version 4.5.1).

## Supporting Information

Supporting Information is available from the Wiley Online Library.

## Acknowledgement

This work was supported by the Stanford Bio-X Interdisciplinary Initiatives Seed Grants Program (IIP) [R10-14], Stanford Center for Cancer Nanotechnology Excellence for Translational Diagnostics (CCNE-TD) seed grant, Stanford Beckman Center Technology Development Seed Grants, the National Cancer Institute of the National Institutes of Health (Award Number R21CA261643, R01CA283247). This work was also supported in part by the Center for Cellular Construction, which is a Science and Technology Center funded by the National Science Foundation (NSF Award: DBI-1548297). Device fabrication and characterization were performed in the Stanford Nanofabrication Facilities (SNF) (NSF Award No. ECCS-1542152) and Stanford Nano Shared Facilities (SNSF) (NSF Award No. ECCS-2026822) respectively. This research was performed in part at the Environmental Molecular Sciences Laboratory, a DOE Office of Science User Facility sponsored by the Biological and Environmental Research program under Contract No. DE-AC05-76RL01830. We thank Marina Dolivo at Google X for assistance with the Nanoscribe Quantum X system and Dr. Matthew Monroe from Pacific Northwest National Laboratory for assistance with MassIVE upload.

## Conflict of Interest

The authors declare no conflict of interest.

## Data Availability Statement

The mass spectrometry raw data have been deposited to the ProteomeXchange Consortium via the MassIVE partner repository with dataset identifiers (MSV000099758, password: Laser7133).

## Table of Contents

Multi-tiered μDicers mechanically dissect tissue slices into uniform microtissues as small as 10 μm. The hierarchical blade architecture limits instantaneous blade-tissue engagement and lowers the cutting force relative to single-tier designs. When coupled with ultrasensitive mass spectrometry workflows, μDicers reveal deeper proteome coverage at cellular scales than laser capture microdissection, offering a powerful tool for next-generation spatial proteomics.

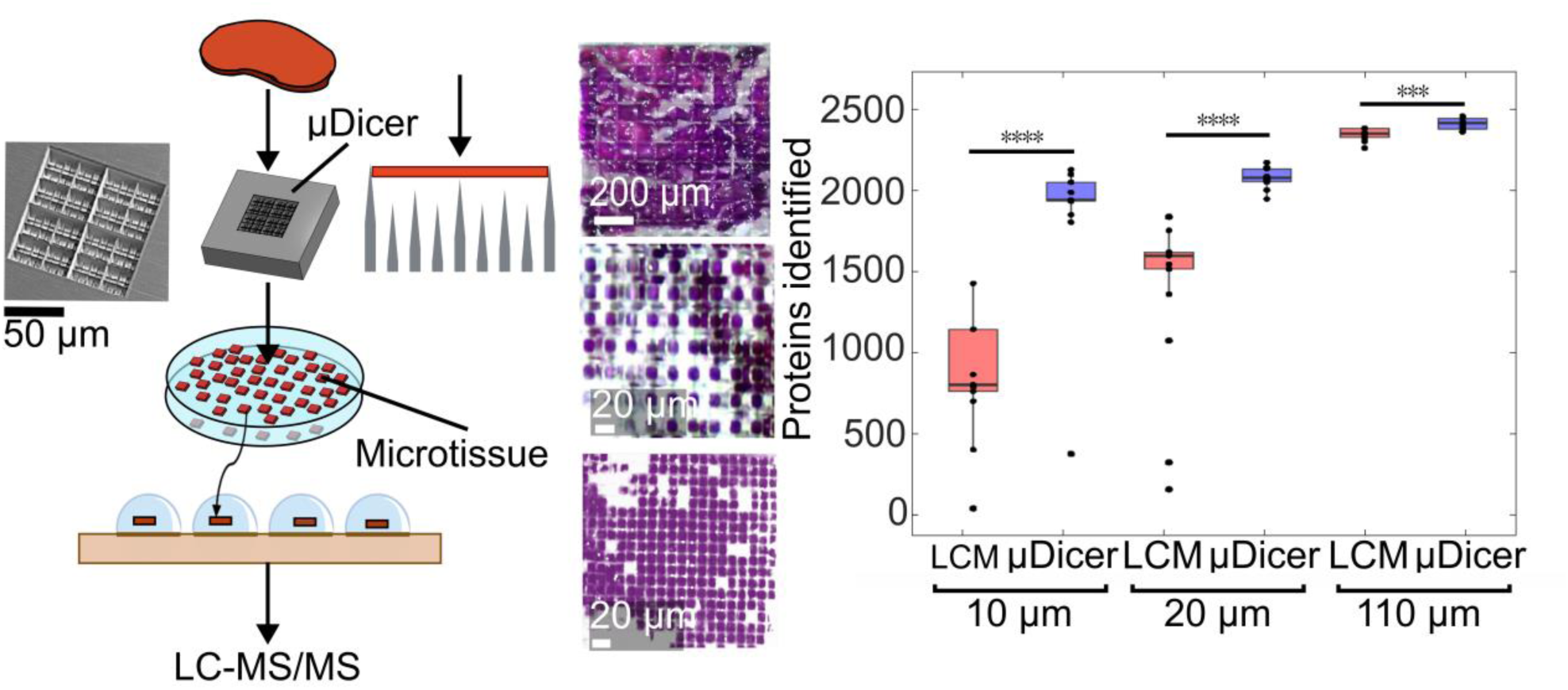

## Notes

### Competing Interest Statement

The authors have declared no competing interest.

