## Supporting figures (S1-S16) and supporting table S1 for "A multi-tiered μDicer with hierarchical blades achieves protein-preserving microdissection down to 10 μm"

### Supporting Information

**Figure S1.** Scheme showing the microdissection of a tissue slice in our multi-tiered μDicer. Section b-b illustrates the idealized dicing process at consecutive time points i.e., $t_0$, $t_1$, $t_2$, and $t_3$ for generating microtissues utilizing a 4-tiered μDicer.

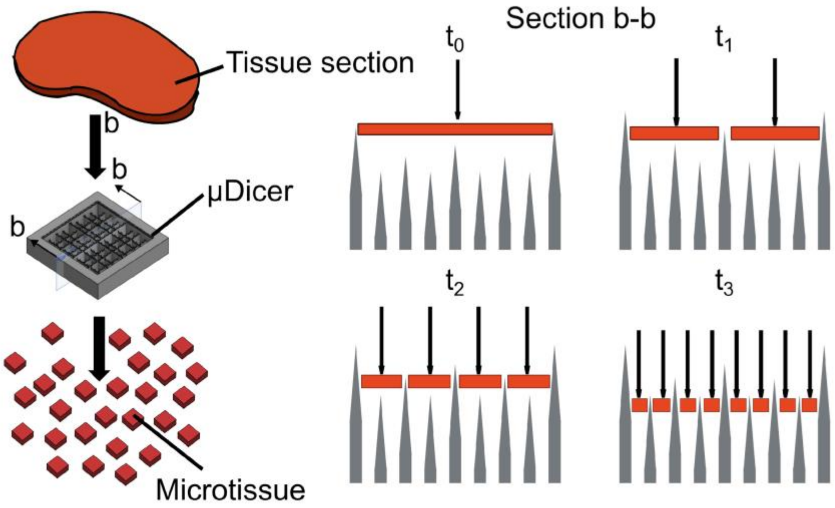

**Figure S2.** We used two-photon polymerization (2PP) additive manufacturing technology to fabricate the µDicers. We printed our µDicers with the Nanoscribe Photonic Professional GT or Quantum X shape using a 25x objective in IP-S photopolymer resin on an indium tin oxide (ITO) coated glass substrate.

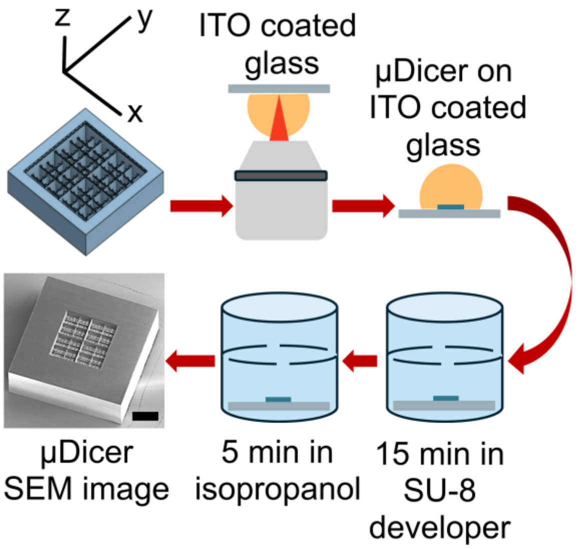

**Figure S3.** CAD (left panels) and scanning electron microscope (SEM) images (right panels) showing μDicers with blade tip spacing of 35 μm in the x-y direction. a) Single-tiered μDicer with serration. b) 2-tiered μDicer with serration. c) 4-tiered μDicer with serration.

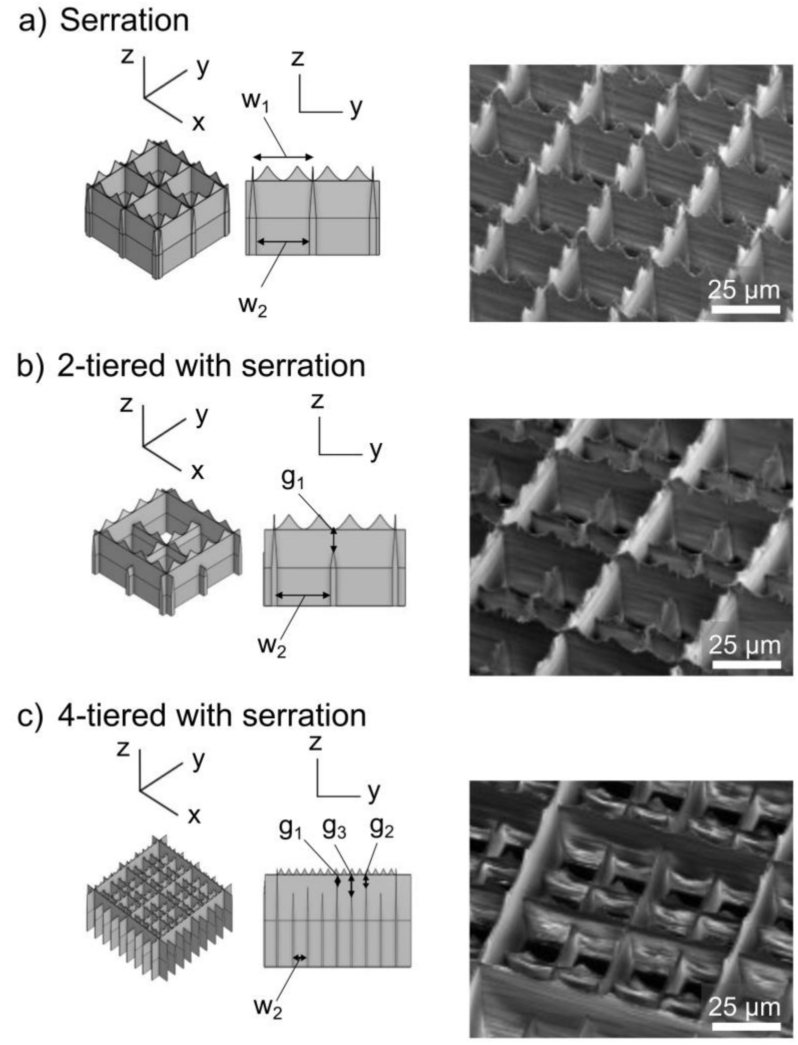


| Symbol | Measurement | Design value |
|---|---|---|
| $w_1$ | Blade tip spacing | 35 μm |
| $w_2$ | Blade base gap width | 25 μm |
| $g_1$ | Reduction in blade base height | 20 μm |
| $g_2$ | Reduction in blade base height | 40 μm |
| $g_3$ | Reduction in blade base height | 60 μm |

**Figure S4.** CAD (left panels) and scanning electron microscope (SEM) images (right panels) showing μDicers with blade tip spacing of 20 μm the in x-y direction. a) Single-tiered μDicer with serration. b) 2-tiered μDicer with serration. c) 4-tiered μDicer with serration.

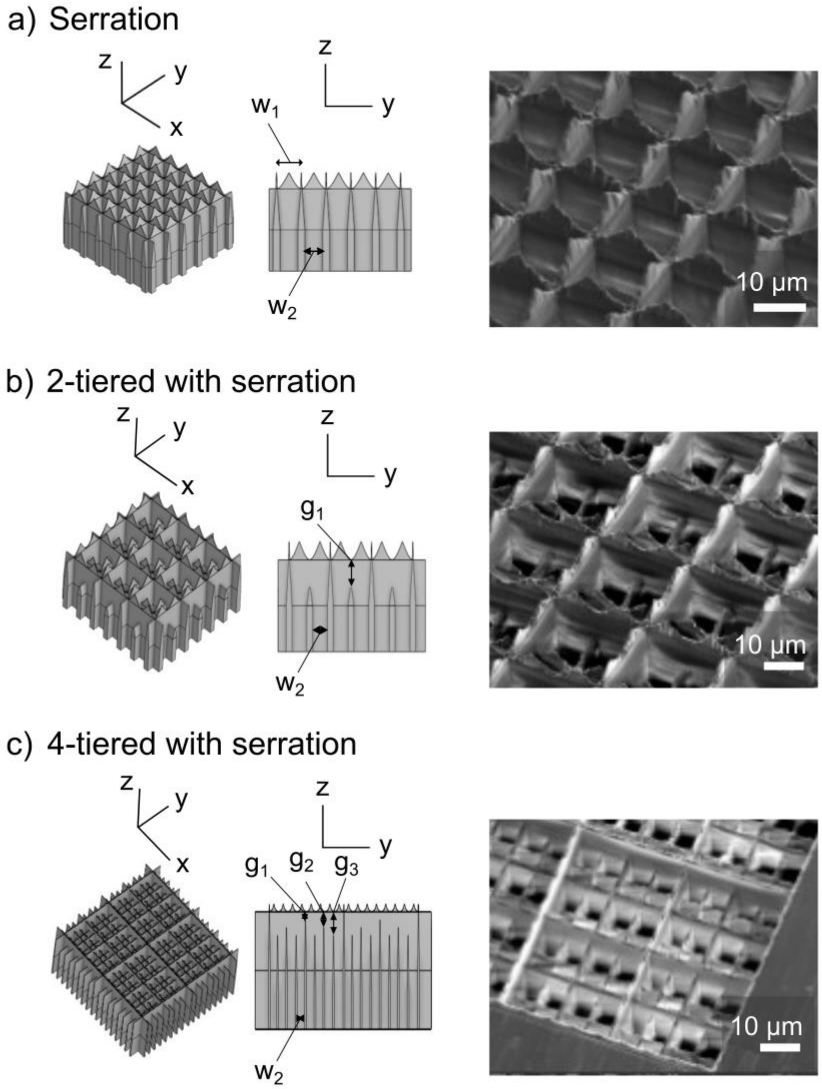


| Symbol | Measurement | Design value |
|---|---|---|
| $w_1$ | Blade tip spacing | 20 μm |
| $w_2$ | Blade base gap width | 10 μm |
| $g_1$ | Reduction in blade base height | 20 μm |
| $g_2$ | Reduction in blade base height | 40 μm |
| $g_3$ | Reduction in blade base height | 60 μm |

**Figure S5.** CAD (left panels) and scanning electron microscope (SEM) images (right panels) showing μDicers with blade tip spacing of 10 μm in the x-y direction. As the blade base width is 10 μm, there is no through-holes in these μDicers. a) Single-tiered μDicer with serration. b) 2-tiered μDicer with serration. c) 4-tiered μDicer with serration.

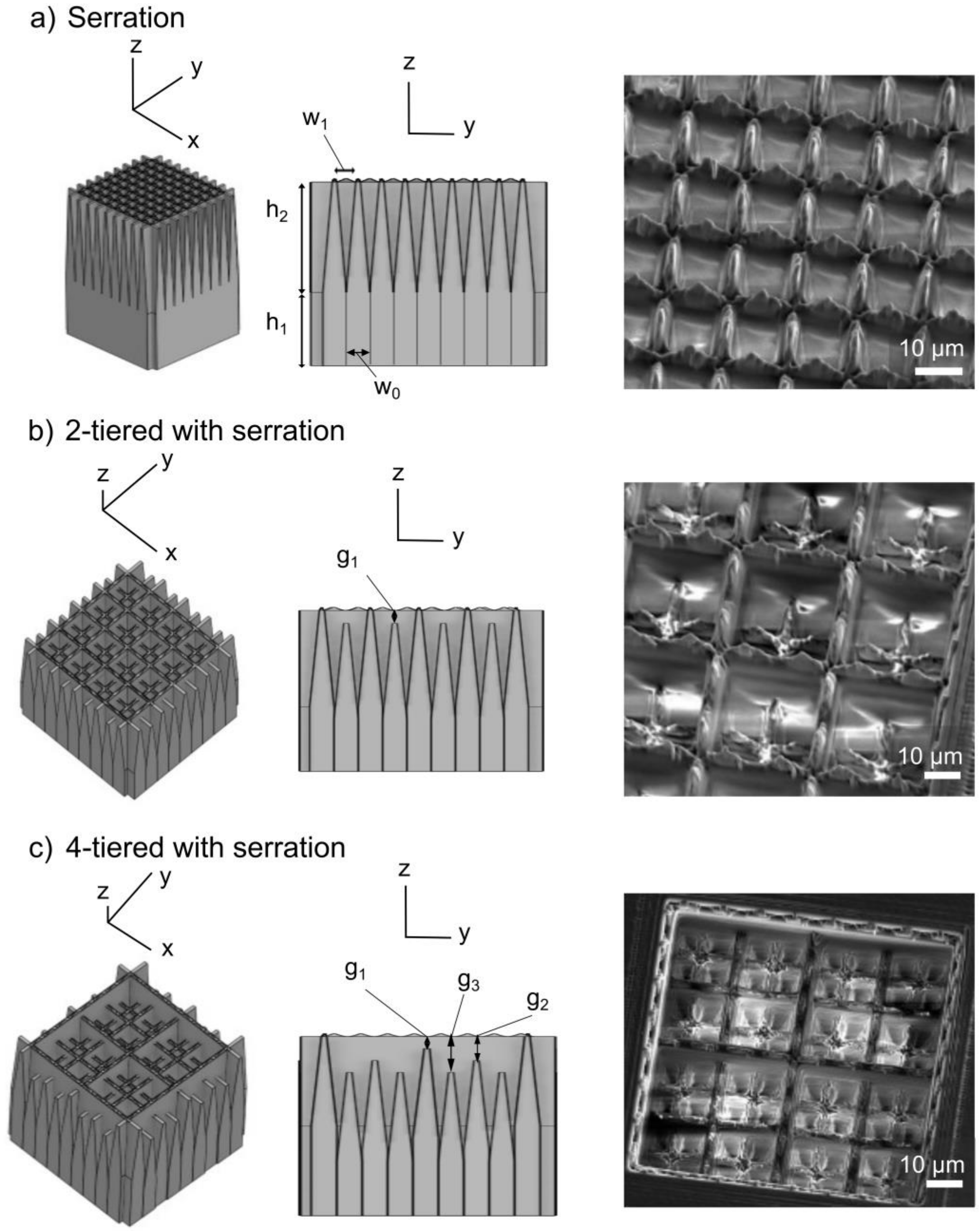


| Symbol | Measurement | Design value |
|---|---|---|
| $w_0$ | Blade base width | 10 µm |
| $w_1$ | Blade tip spacing | 10 µm |
| $h_1$ | Blade base height | 150 µm |
| $h_2$ | Blade tip height | 150 µm |
| $g_1$ | Reduction in blade base height | 20 µm |
| $g_2$ | Reduction in blade base height | 40 µm |
| $g_3$ | Reduction in blade base height | 60 µm |

**Figure S6.** a) Images show "microtissues" generated from various samples: agar (2%) (1.1 and 2.1), wax (1.2 and 2.2), fixed beef liver (1.3 and 2.3), and fixed LTL610, a human prostate cancer tissue grown in a mouse (1.4 and 2.4). Microtissues were generated using µDicers with flat blades and 2-tiered blades with serration. The blade tip spacing was 110 µm. Prior to dicing, all samples were pre-sliced to a thickness of approximately 0.2 to 1 mm using Compresstome® tissue slicer. Both beef liver and LTL610 were fixed using 50% ethanol (30 seconds), 70% ethanol (30 seconds), 90% ethanol (30 seconds), and 95% ethanol (30 seconds) respectively. All ethanol solutions were pre-chilled at 4 °C before fixation which occurred at room temperature. b) Box plot presents microtissue widths obtained from image analysis in ImageJ. Widths were determined by measuring the narrowest dimension passing through the center of each microtissue. Sample sizes were n = 51 for agar (2%) and wax, and n = 31 for beef liver and LTL610. Median widths are indicated by red horizontal lines.

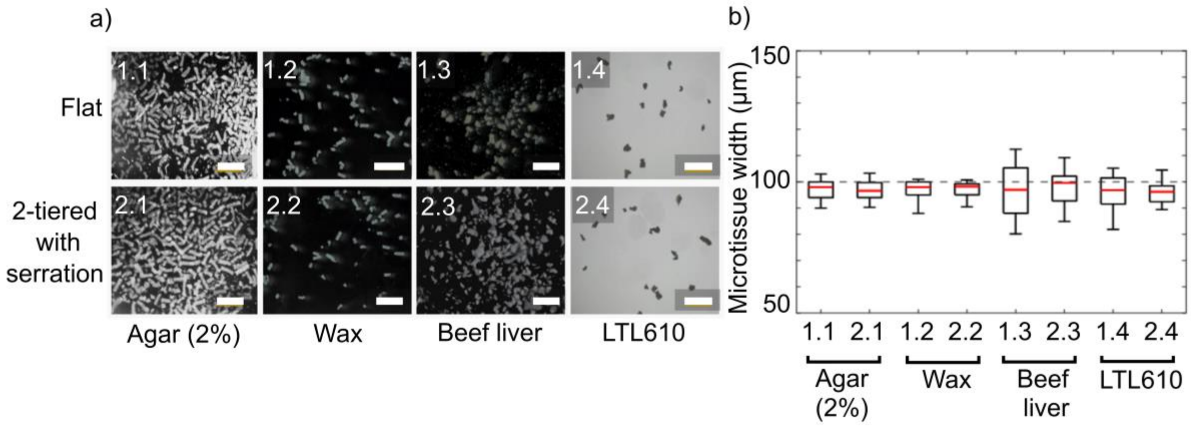

**Figure S7.** Force-displacement curves for 2-tiered and 4-tiered µDicers with blade tip spacings indicated in the graph (i.e., 10 µm, 20 µm, 35 µm, and 110 µm). Each data point was measured at a different location on the same SCC tissue slice (organ: soft tissue). For 2-tiered and 4-tiered µDicers with 10 µm blade tip spacing, we used SCC tissue slice derived from lymph node to collect the data. Note that the µDicers with different blade tip spacings had different blade array area. For the µDicers with 10 µm, 20 µm, 35 µm, and 110 µm blade tip spacings, the blade grid array areas were 80 µm x 80 µm, 160 µm x 160 µm, 200 µm x 200 µm, and 1 mm x 1 mm, respectively. These designs still allowed us to evaluate the effect of different blade geometries for a given blade tip spacing while reducing print time in the Nanoscribe.

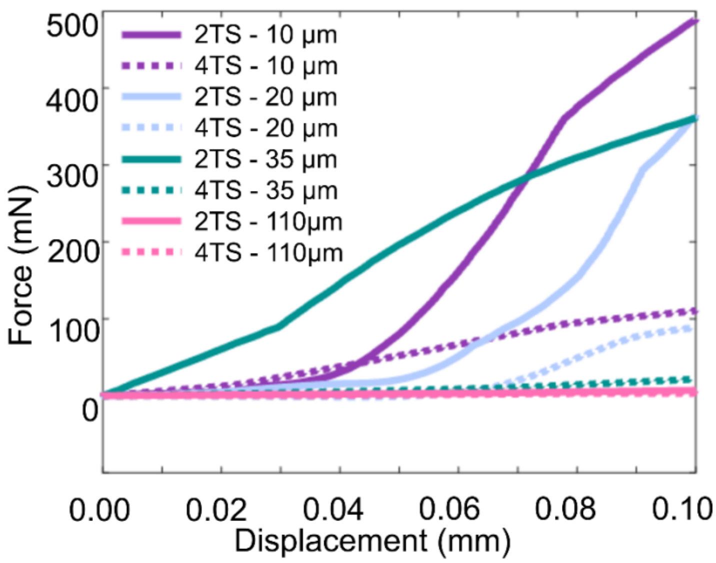


| Device | ID |
|---|---|
| 2-tiered with serration | 2TS |
| 4-tiered with serration | 4TS |

**Figure S8.** a) Reusability assessment of a single flat blade. SEM images of the blade before and after one and two cuts, as well as the cut mark on the hematoxylin and eosin (H&E) stained, ethanol-fixed squamous cell carcinoma (SCC) slice. b) Force displacement analysis of the first and the second cut. Increased force was required for the second cut (in a new region on the tissue slice), indicating the blunting of the blade tip.

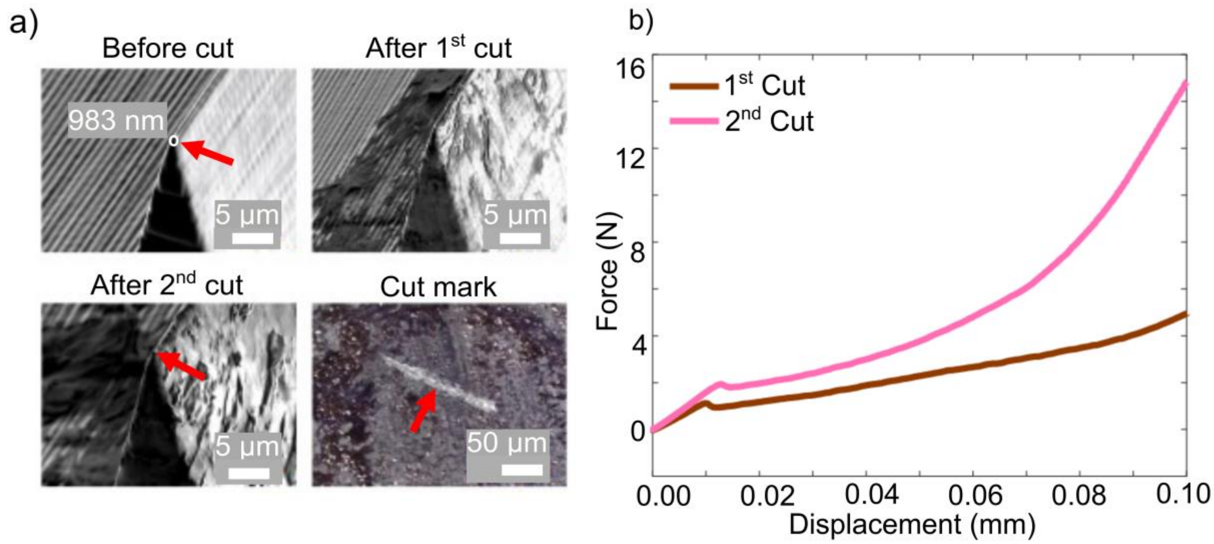

**Figure S9.** a-c) Images of hematoxylin and eosin (H&E) stained, ethanol-fixed squamous cell carcinoma (SCC) tissue slices (10 µm thick) after being cut by a tissue chopper (McIlwain) with a stainless-steel razor blade at 100 µm, 25 µm, and 10 µm displacements, respectively. The cutting was irregular and ineffective, likely due to excess heat generation during the cutting process. Tissue cutting in such small dimensions also resulted in low microtissue collection efficiency. We were only able to collect 2 out of 100 microtissues due to the scattering of the cut microtissues with 100 µm displacement. In this experiment, the tissue slices were placed on poly-L-lysine-coated glass slides. We attempted placing the tissue slice on a wax-coated glass slide, but the tissue was not cut due to poor adhesion between the tissue slice and wax, possibly arising from the heat generation during the cutting process. d) Magnified image of one microtissue generated by the tissue chopper with 100 µm displacement.

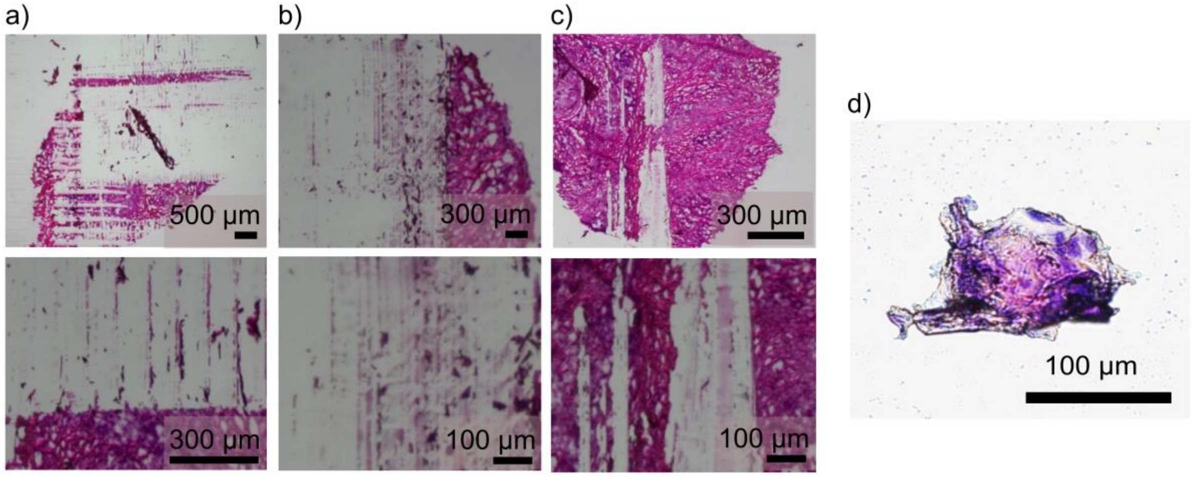

**Figure S10.** Number of a) proteins and b) peptides identified by tandem mass spectra (MS2) (i.e., without match-between-run) from squamous cell carcinoma (SCC) microtissues of different sizes, generated using LCM and μDicers. Wilcoxon rank-sum test was used for the comparisons between LCM vs. μDicers: * $p \leq 0.05$; ** $p \leq 0.01$; *** $p \leq 0.001$; **** $p \leq 0.0001$.

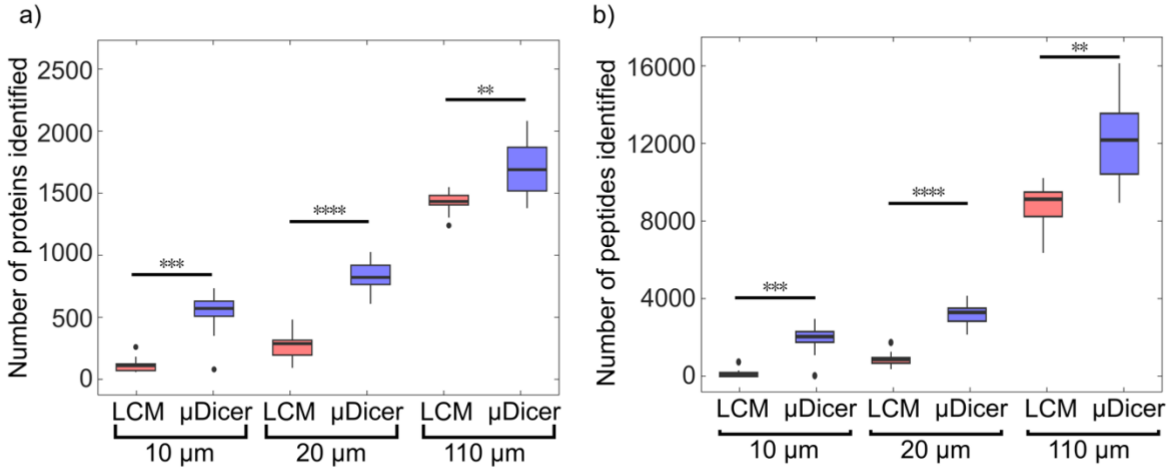

**Figure S11.** a) Percentage of peptide-spectrum matches (PSMs) for full-tryptic, semi-tryptic, and non-tryptic peptides, and b) methionine-oxidized peptides from microtissues generated by LCM and μDicer at 110 μm lateral resolution. Comparison of LCM vs. μDicer yielded *p* values of 0.030, 0.028, 0.62 for full-tryptic, semi-tryptic, and non-tryptic peptides, respectively; and 0.27 for methionine-oxidized peptides.

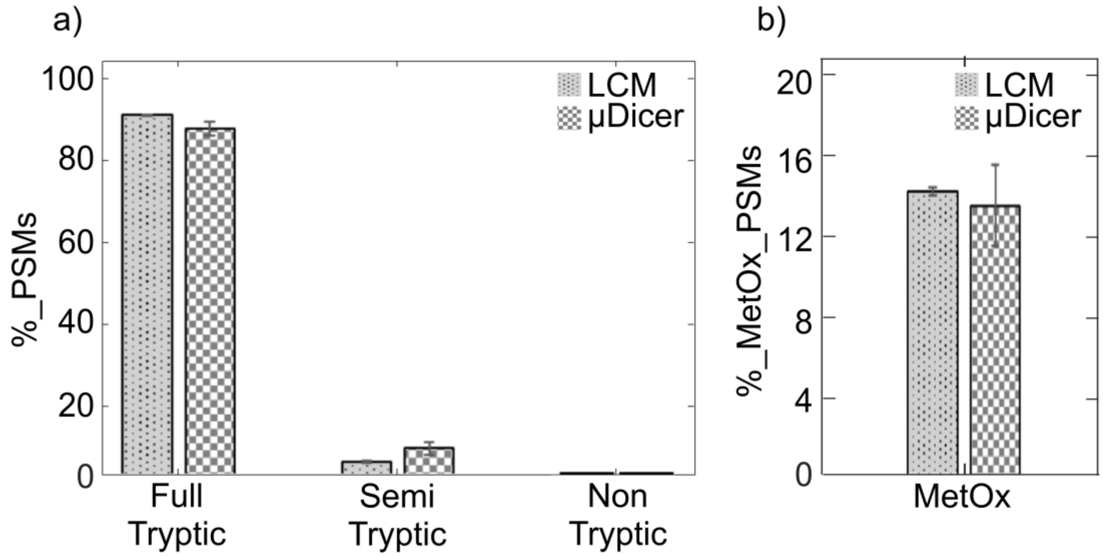

**Figure S12.** Pearson correlation coefficients of protein intensities calculated for microtissues of various sizes, generated by LCM or µDicer. Intensities of proteins identified from µDicer-generated microtissues showed higher correlation than LCM-generated microtissues across microtissue sizes.

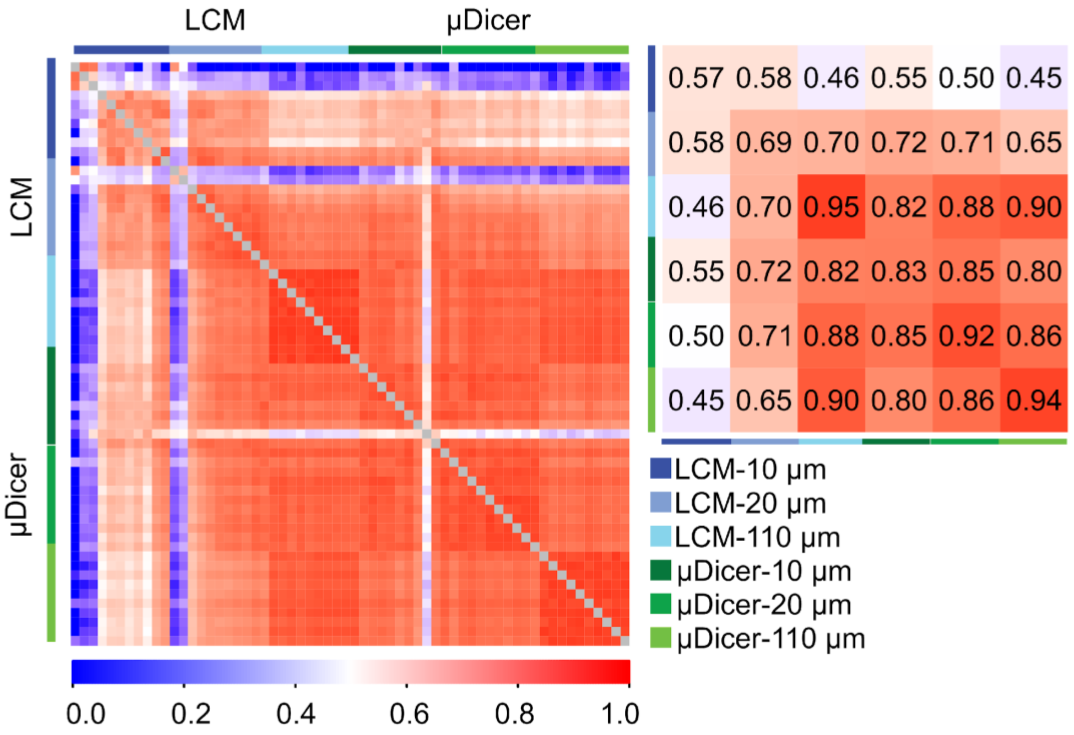

**Figure S13.** Venn diagrams showing the overlap of proteins detected from microtissues of various sizes generated using a) LCM and b) µDicer. c) Venn diagram showing the overlap of proteins detected from 10 µm microtissues generated using LCM or µDicer.

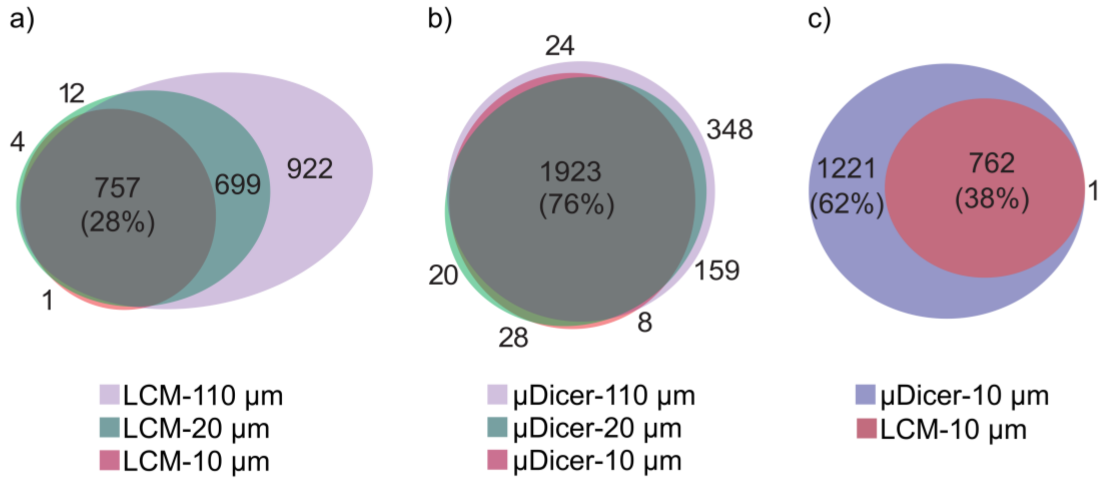

**Figure S14.** Characterization of the actual size of PEN membrane and microtissues (adhered to PEN membrane) after cutting and catapulting using LCM. a) Brightfield images and b) 3D rendering of confocal fluorescence z-stacks of blank PEN membranes (with no tissue slices mounted) reveal deformation at the center of the samples, with red arrowheads marking dark spots at these regions following cutting and catapulting by LCM. c) Brightfield images and b) 3D rendering of confocal fluorescence z-stacks of microtissues (adhered to PEN membrane) exhibit similar deformation and material loss as observed in the blank PEN membrane samples. Red arrowheads in d) highlight through-holes and areas of significant material loss. The volumes of the microtissues shown with nominal sizes of 110 μm, 20 μm, and 10 μm are measured to be 61956 $\mu m^3$, 1925 $\mu m^3$, and 253 $\mu m^3$ respectively, corresponding to only 51%, 48%, and 25% of their expected volumes.

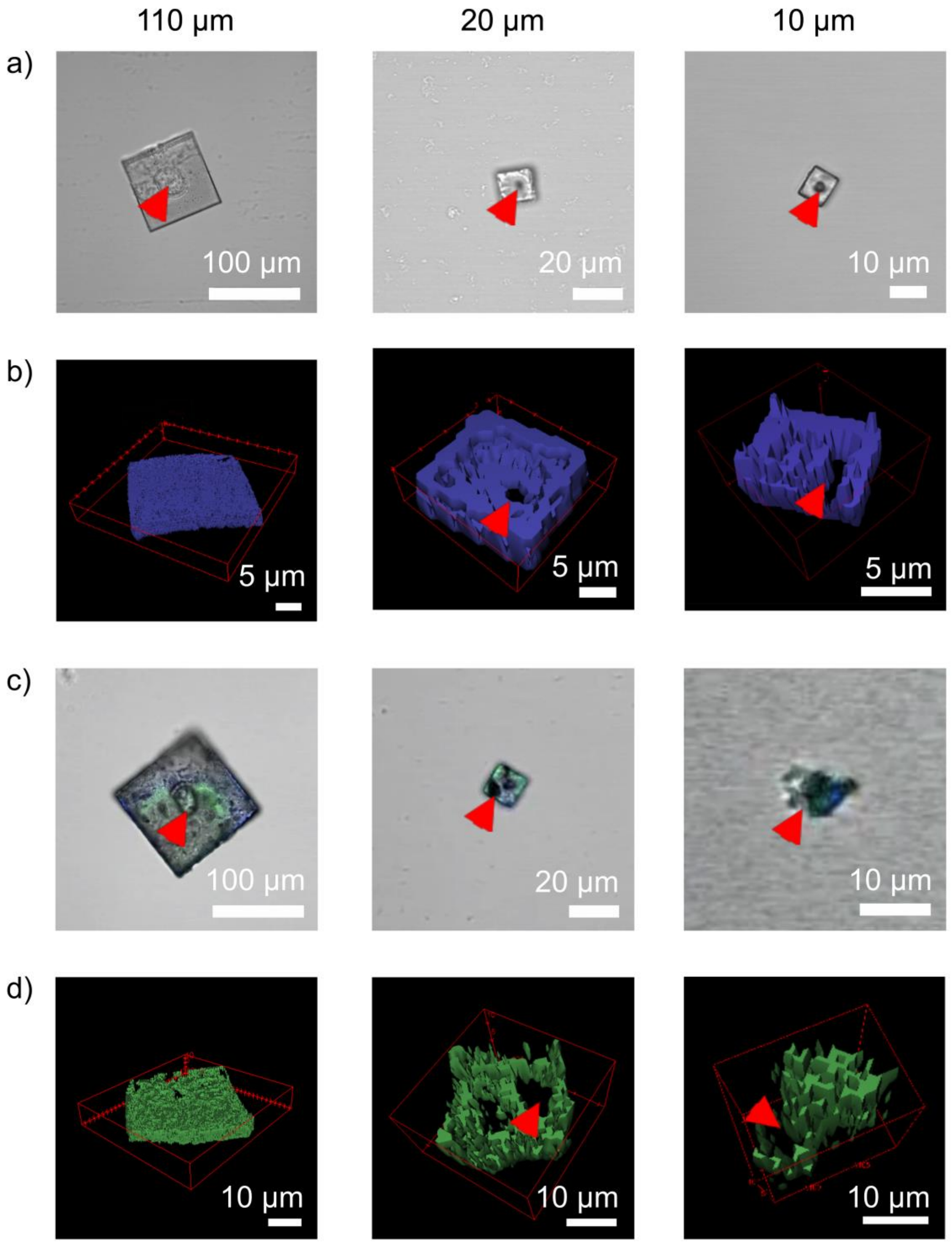
110 µm
20 µm
10 µm
a)
100 µm
20 µm
10 µm
b)
5 µm
5 µm
5 µm
c)
100 µm
20 µm
10 µm
d)
10 µm
10 µm
10 µm

**Figure S15.** Atomic force microscopy (AFM) measurements of tissue stiffness. a) AFM cantilevers on top of a hematoxylin and eosin (H&E) stained, ethanol-fixed squamous cell carcinoma (SCC) tissue slice (obtained from lymph node) in 70% ethanol. Tissue slices (100 µm thick) were sectioned using a cryostat and placed on top of glass slides coated with 0.1 mg/mL poly-L-lysine (incubated overnight and rinsed with DI water). The tissue slices were then fixed with ethanol and stained using H&E staining medium. The AFM indentation measurements were conducted using a cantilever with a 5 µm tip radius, 150 µm length, 18 µm width, 16.0 kHz frequency, and a spring constant of 0.115 N/m. The SCC tissue slice exhibited heterogeneity across the scanned region (~ 40 µm across). Four data points were collected per scanned region. b) The box plot summarizes stiffness measurements from 55 locations on the tissue slice. The values ranged from 19.4 kPa to 602.6 kPa, which indicates heterogeneity in the SCC sample. The mean stiffness was 155.4 kPa, while the median was 103.2 kPa.

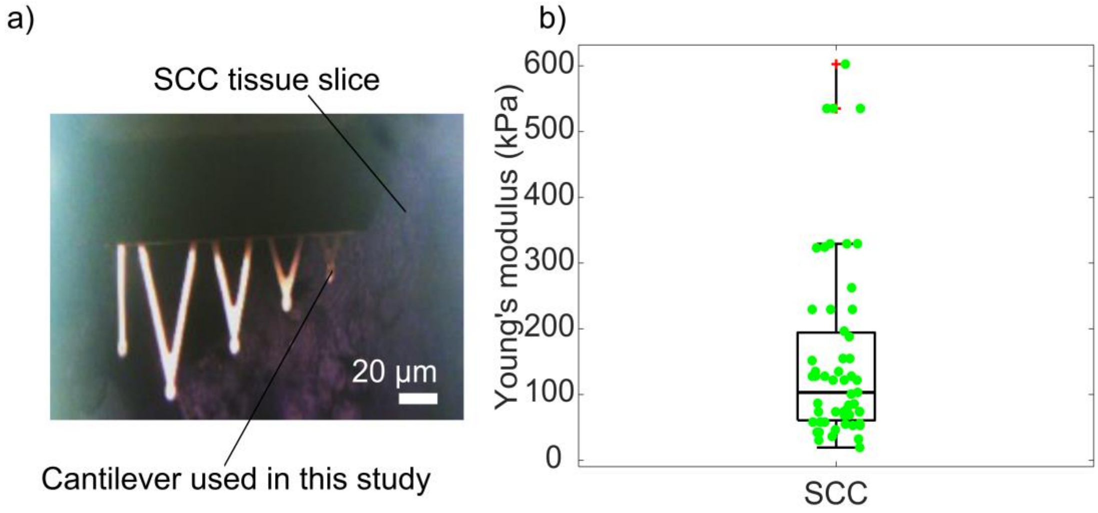

**Figure S16.** a) Top view of a 4-tiered µDicer with 110 µm blade tip spacing, with a stained and ethanol-fixed squamous cell carcinoma (SCC) tissue slice placed on top. b) Top view of the same µDicer following the release of SCC microtissues. No microtissue remained inside the µDicer after the release protocol. c) Top view of a 4-tiered µDicer with 10 µm blade tip spacing after dicing an SCC tissue slice but before release. d) Top view of the same µDicer after releasing the diced microtissues.

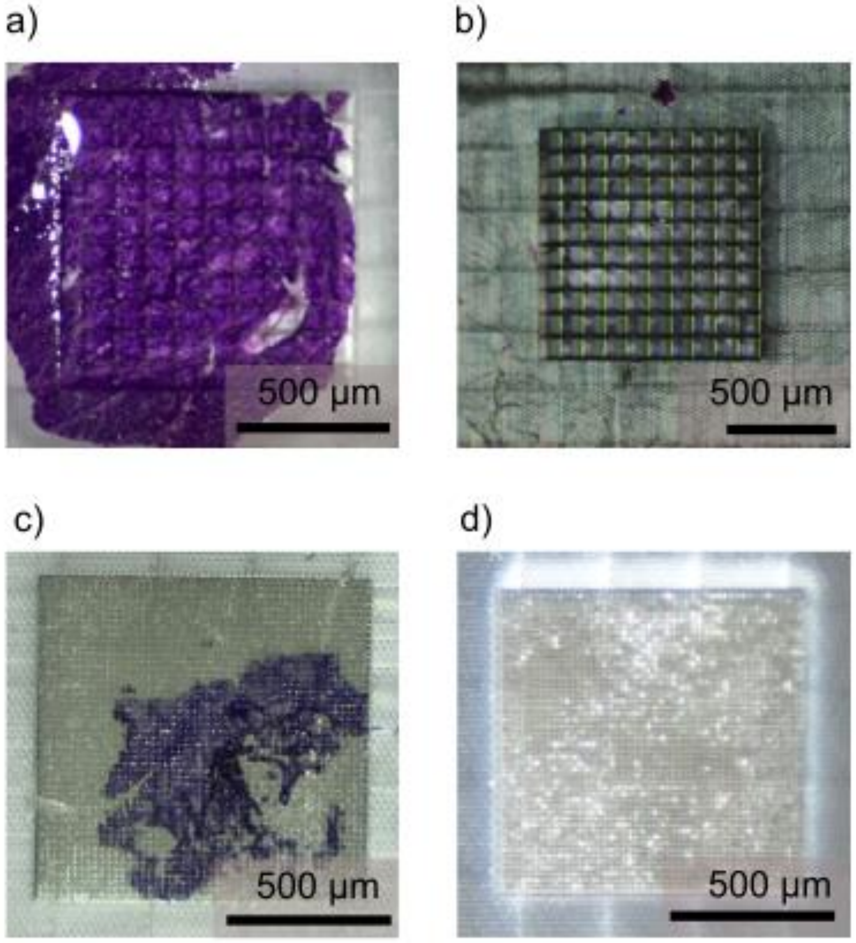

**Table S1:** Measurements of SCC microtissue volumes generated by LCM and µDicer at 10 µm and 20 µm lateral resolutions. To be consistent with other presented data in this work, the LCM-generated microtissues were collected at PNNL and imaged at Stanford using confocal microscope (Zeiss LSM-780). The volume was calculated from confocal fluorescence images of microtissues generated using LCM and µDicer (see Experimental Methods). For LCM-generated 20 µm microtissues, 8 samples were shipped in nanoPOTS chip from PNNL and all those 8 samples were analyzed. For LCM-generated 10 µm microtissues, 14 samples were shipped in nanoPOTS chip. However, only 9 samples were found in the wells of nanoPOTs chip for confocal imaging. Other wells containing 10 µm microtissues were empty, likely due to sample loss during shipping. Of the 9 samples, only three contained identifiable SCC microtissues and were analyzed; the remaining six contained debris, PEN membrane only, or were lost during manual transfer. For µDicer-generated microtissues, 10 samples of each size were analyzed.

| Expected microtissue size | 20 x 20 x 10 $\mu m^3$ (4000 $\mu m^3$) | | 10 x 10 x 10 $\mu m^3$ (1000 $\mu m^3$) | |
|---|---|---|---|---|
| Device | LCM | µDicer | LCM | µDicer |
| Volume ($\mu m^3$) | 1925 | 4290 | 253 | 1033 |
| | 2138 | 4019 | 101 | 858 |
| | 2269 | 4069 | 889 | 817 |
| | 2227 | 3520 | - | 1005 |
| | 3055 | 4072 | - | 979 |
| | 2537 | 3958 | - | 922 |
| | 1949 | 3899 | - | 1003 |
| | 1433 | 4014 | - | 998 |
| | - | 3681 | - | 962 |
| | - | 3915 | - | 1024 |
| Mean ($\mu m^3$) | 2191.6 | 3943.7 | 414.3 | 960.1 |
| % of expected volume | 54.8 | 98.6 | 41.4 | 96.0 |
| CV (%) | 21.7 | 5.4 | 100.9 | 7.6 |
